# The Internal Structure of Seizures Reflects Animal-Specific Trajectories of Disease Progression and Circadian Modulation in Epileptic Mice

**DOI:** 10.64898/2026.08.21.746213

**Authors:** Vinicius Lima, Antoine Ghestem, Viktor Jirsa, Christophe Bernard, Damien Depannemaecker

**Affiliations:** Aix Marseille Université, INSERM, INS, Institut de Neurosciences des Systèmes, Marseille, France; Institut de Neurosciences de La Timone, UMR 7289, CNRS, Aix-Marseille Université, Marseille 13005, France

## Abstract

Epileptic seizures arise from the abnormal, sustained recruitment of neuronal populations, yet the temporal organisation of activity *within* individual seizures remains poorly understood. We hypothesised that seizures follow structured trajectories through a restricted state space rather than random patterns of activity. To test this idea, we analysed continuous long-term EEG recordings from six mice with pilocarpine-induced temporal lobe epilepsy and developed a symbolic framework that transforms seizure activity into sequences of discrete burst states. Across more than one thousand spontaneous seizures, we found that seizures are organised by sparse, animal-specific transition rules linking a small repertoire of recurrent burst motifs. These rules evolve over the course of epilepsy: early seizures display substantial variability in burst composition, and, to a smaller but statistically significant degree, in transition structure, whereas later seizures become more stereotyped in which burst types they use, a pattern suggestive of consolidation of the epileptic network into a stable dynamical regime. Seizure microstructure was further modulated by circadian phase, indicating that biological rhythms influence not only when seizures occur but also how they unfold. Moreover, burst sequences exhibited higher-order temporal dependencies that could not be explained by first-order Markov statistics alone. Together, these results reveal an evolving seizure grammar that links ictal dynamics to disease progression and circadian regulation, establishing symbolic dynamics as a powerful framework for quantifying the internal organisation of seizures.

**significance statement:** Why seizures differ from one another, and how those differences relate to the evolution of epilepsy in each subject, remain poorly understood. We demonstrate that seizures are structured dynamical trajectories rather than uniform episodes of abnormal activity. Using symbolic and information-theoretic analyses of long-term recordings in epileptic mice, we identify a limited repertoire of recurrent activity patterns whose organisation evolves during chronic epilepsy and is modulated by circadian phase. These findings suggest that the internal structure of seizures encodes information about network reorganisation over time, providing a class of biomarkers for tracking disease progression.

## I. INTRODUCTION

Epileptic seizures are characterised by sudden, abnormal electrical activity in the brain, often leading to significant neurological and physiological consequences [7]. Extensive research has focused on the dynamical properties at seizure onset, in particular the identification of pre-ictal biomarkers that could enable seizure prediction [16, 32]. Complementary work on seizure dynamotypes has shown that the bifurcation mechanism governing seizure onset and offset is not fixed within an individual, but can itself evolve over the course of epileptogenesis and chronic disease [29], underscoring that seizure dynamics are non-stationary at the level of onset mechanism as well as internal structure. By contrast, the temporal structure within individual seizures remains much less characterised. Individual seizures contain sequences of morphologically distinct bursts of activity whose ordering, recurrence, and transition statistics could, in principle, reflect the trajectory of a high-dimensional neural system through a constrained state space [3, 5, 11, 18, 28]. Mapping this internal grammar may offer a window into how the network reorganises as epilepsy becomes chronic. Addressing this question requires an experimental model in which seizure activity can be monitored continuously throughout the development of epilepsy. The pilocarpine-induced status epilepticus (SE) model is one of the most widely used models of temporal lobe epilepsy (TLE) in rodents [10, 33]. Following pilocarpine administration, animals undergo an initial episode of SE, a latent period, and subsequently enter a chronic epileptic phase characterised by spontaneous recurrent seizures. In susceptible strains such as FVB mice, long-term telemetric electroencephalographic (EEG) recordings provide dense sampling of seizure activity over weeks to months, enabling the progressive evolution of ictal dynamics to be quantified across the course of epileptogenesis and chronic disease. Seizures in epilepsy do not occur randomly in time. Both circadian and multiday (multidien) rhythms strongly modulate seizure probability [2, 19]. Whether circadian phase also modulates the *internal* structure of individual seizures—that is, whether seizures occurring at different times of day follow distinct dynamical trajectories through seizure state space—has received comparatively little attention. Characterising such a relationship would bridge two largely separate perspectives on epilepsy: the temporal organisation of seizure occurrence across days and the dynamical organisation of activity within individual seizures.

The temporal structure of complex signals is naturally captured by symbolic dynamics [26]. Converting a continuous time series to a discrete sequence of symbols allows the application of tools from information theory and stochastic processes: Shannon entropy quantifies diversity, Markov transition matrices capture sequential dependencies, Kullback-Leibler (KL) divergence measures the similarity between distributions, and block-entropy analysis estimates the effective memory length of the sequence [24, 31]. These methods have been applied to neural signals to study brain state transitions [4], microstate sequences in EEG [26], and, more recently, to characterise information processing in epileptic circuits [7, 22].

Here we build directly on these frameworks. We define dynamical states as distinct burst types detected in the LFP and clustered by their morphological features, then encode each seizure as a symbolic sequence of burst-type labels. We ask four related questions: (i) What are the transition statistics between burst types within seizures? (ii) Does the transition structure change over the course of the recording, and if so, when? (iii) Is the entropy of burst-type occupancy modulated by time of day? (iv) Do burst sequences carry higher-order (beyond first-order Markov) temporal dependencies, and does this change with disease progression?

## II. METHODS

### 1. Animals and housing

Six FVB adult male mice (Charles River) were used in this study. The animals were maintained on a 12-hour light/dark cycle (7 am–7 pm light), with ad libitum access to food and water. Continuous electroencephalography (EEG) monitoring was achieved using telemetric probes (ETA-F10, Data Sciences International). All procedures were performed in accordance with institutional animal care guidelines.

### 2. Surgical Procedure

Surgical procedures were performed under anaesthesia using a combination of Xylazine (10 mg/kg) and Ketamine (100 mg/kg), with additional local analgesia provided by Bupivacaine. Telemetric probes were implanted subcutaneously in the back of the animals, and two wires were connected to microscrews (000-120, stainless steel). One recording screw was placed above the dorsal CA1 (stereotaxic coordinates: anteroposterior: −1.8 mm; mediolateral: −1.8 mm), and the other was positioned above the cerebellum to serve as a reference. The screws were covered with dental cement (Paladur, Kulzer) for electrical insulation. Surgical procedures were performed 10 days prior to pilocarpine injection.

### 3. Model of epilepsy

The mice were pre-treated with methylscopolamine (1 mg/kg, i.p., Sigma Aldrich) 30 minutes before the pilocarpine injection to minimise peripheral effects. Pilocarpine was repeatedly injected (100 mg/kg, i.p., Sigma Aldrich) every 20 minutes until Status Epilepticus (SE) was established. After 90min of SE, we injected diazepam (10 mg/kg, i.p., Diazedor, Axience) to stop SE.

### 4. EEG Recording and Data Acquisition

EEG signals were recorded continuously using a DataQuest Acquisition System ART 4.1 (Data Sciences International), with a sampling rate of 500 Hz.

### 5. Data Analysis

Seizure events were semi-automatically detected using Neuroscore software (Data Sciences International). Detection was based on predefined threshold values (spike amplitude, minimum/maximum spike duration, interval between spikes, and minimum number of spikes).

### 6. Seizure snippet extraction

Detected seizures were extracted from the continuous EEG by aligning each seizure to its onset time and retaining a window extending from *w*_prev_ = 30 seconds before onset to *w*_aft_ = 30 seconds after the end of the longest seizure in each animal. This ensures that all seizures are time-aligned and have a common duration.

### 7. Detection of bursts and dynamical states

The full pipeline, from raw LFP to a discrete symbolic sequence of burst states, proceeds in four stages: adaptive segmentation, feature extraction, unsupervised clustering, and state-vector construction. Each stage is described below and illustrated in Figure 1.

**Figure 1.**
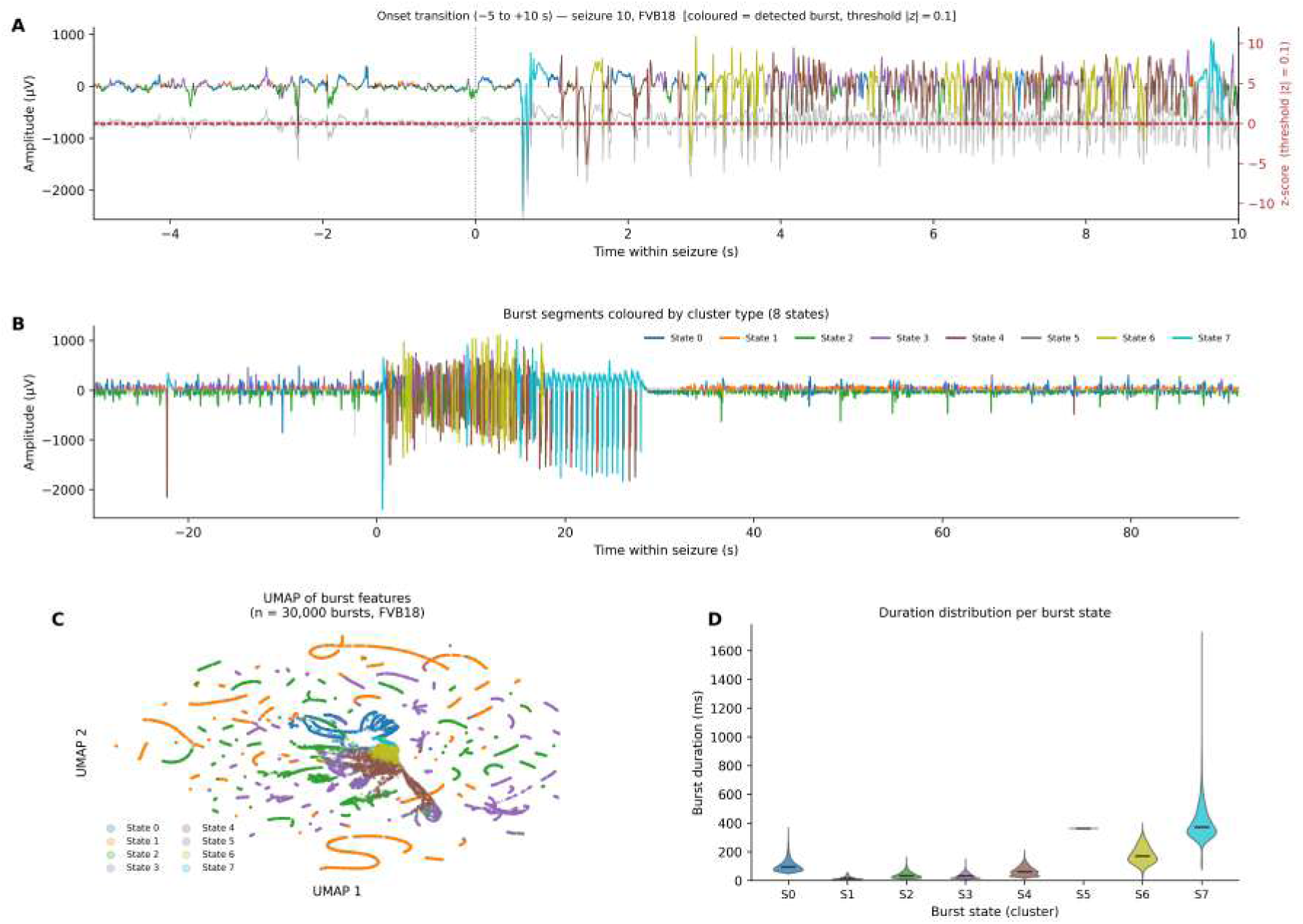
Burst detection, feature extraction, and unsupervised state clustering pipeline (FVB18, seizure 10). **(A)** Fifteen-second window spanning the seizure onset (*t* = 0, dotted vertical line). The grey trace shows the raw LFP amplitude (left axis); the lighter grey trace on the right axis shows the locally z-scored signal (Eq. 1) computed within the 30-second window that contains this segment, with the detection threshold |*z*| = 0.1 indicated by red dashed lines. Burst segments, defined as contiguous runs of samples with |*z*| ≥ 0.1, are coloured by their assigned cluster state (same palette as panels B–D). During the pre-onset period (left) the signal is predominantly near baseline with only sparse, low-amplitude burst segments; after onset the trace is dominated by burst activity of diverse morphology and cluster identity. A brief interval of residual background can appear immediately after the annotated onset (here, ∼ 0.5 s), reflecting the short lag between the marked onset time and the first threshold-crossing burst rather than a segmentation failure. The |*z*| = 0.1 threshold is intentionally permissive: nearly all oscillatory activity is labelled, and uncoloured (background, −1) regions correspond only to the narrow band of samples within 0.1 standard deviations of the local window mean. **(B)** Full seizure trace (−25 to +90 s relative to onset). Burst segments are coloured by cluster label (states 0–7); background activity is grey. A clear shift in dominant colour is visible across the seizure: the early ictal period (0–30 s) is dominated by high-amplitude bursts from states 2 and 4 (cyan/green), whereas the later ictal and post-ictal periods contain lower-amplitude, morphologically diverse bursts from states 0, 1, and 7 (red/orange tones). This temporal organisation of burst-type sequences is the object of the Markov and symbolic analyses in the Results. **(C)** UMAP [21] projection of the 10-dimensional standardised feature space for 30,000 randomly sampled bursts from FVB18. Each point is one burst, coloured by its *k*-means cluster label. The eight states occupy largely non-overlapping regions of the manifold, confirming that the clustering captures morphologically distinct burst populations. The ribbon-like manifold topology suggests that burst morphology varies principally along a small number of continuous axes. **(D)** Violin plots of burst duration (ms) per cluster state. States S1 and S3 are dominated by brief bursts (*<* 50 ms), consistent with rapid, high-frequency oscillatory events in the early ictal period, whereas state S6 exhibits a broad distribution extending up to ∼ 280 ms and state S7 is concentrated at even longer durations, extending beyond 250 ms, reflecting sustained low-frequency discharge events. These systematic differences in duration alone illustrate the morphological separability of the cluster states.

*a. Stage 1: Adaptive z-score segmentation.* Epileptic LFP recordings are highly non-stationary: baseline amplitude drifts substantially between seizures, between animals, and across weeks of recording. A fixed voltage threshold would therefore be unreliable. Instead, we normalise the LFP locally within non-overlapping 30-second sliding windows. Within each window *w*, the z-scored signal is:

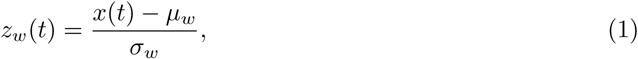

where *µ_w_* and *σ_w_* are the mean and standard deviation of *x*(*t*) computed across all samples in window *w*. This makes the detection threshold adaptive to the local signal statistics, so that the same relative amplitude criterion is applied consistently regardless of absolute amplitude level.

Any contiguous run of samples satisfying |*z_w_*(*t*)| ≥ 0.1 is defined as a *burst segment*. The threshold of 0.1 z-score units is intentionally permissive: it partitions essentially all oscillatory activity into labelled segments, with only the narrow band of samples within 0.1 standard deviations of the local mean remaining as unlabelled *background* (label = −1). This design choice reflects the aim of the pipeline, which is not to distinguish “burst” from “no-burst” in the classical sense, but to segment the full LFP activity stream into morphologically distinct episodes that can then be classified. Adjacent burst segments separated by fewer than two samples are merged into a single event. Detection is done over all seizure trials simultaneously, making it computationally efficient even for recordings spanning hundreds of seizures (Table I).

Figure 1A shows a 15-second window spanning the seizure onset (*t* = 0) for one representative seizure of FVB18. During the pre-onset period (left), activity near the baseline is largely uncoloured (background), while sporadic low-amplitude oscillations begin to appear as the first coloured segments. After onset, burst segments of diverse morphology rapidly dominate the trace, illustrating both the onset of ictal activity and the diversity of burst types (colours) that are recruited within seconds. Because *t* = 0 marks the externally annotated seizure onset rather than the first burst-detection threshold crossing, a brief interval of residual background can persist immediately after onset (here, ∼ 0.5 s) before the first high-amplitude burst is detected; this reflects a short annotation-to-detection lag rather than a failure of the segmentation.

*b. Stage 2: Morphological feature extraction.* Each burst segment is described by a 10-dimensional feature vector. Let *b* = *x*[*t_i_* : *t_f_* ] denote the raw LFP samples of a burst spanning from sample *t_i_* to *t_f_* , and let Δ*t* = *t_f_* − *t_i_* be the duration in samples. The features extracted for each burst are:

1. **Duration** (ms): Δ*t* × 1000*/f_s_*, where *f_s_* = 500 Hz.
2. **Number of peaks** (#peaks): count of local maxima of *b*, detected by scipy.signal.find peaks.
3. **Number of troughs** (#troughs): count of local minima of −*b*.
4. **Dominant frequency** (Hz): (#peaks+#troughs) */* (Δ*t/f_s_*), estimating the mean oscillation frequency from the inter-extremum interval.
5. **Mean voltage** (*µ*): 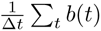.
6. **Voltage standard deviation** (*σ*): std(*b*).
7. **Coefficient of variation** (CV): *σ/*(*µ* + *ɛ*), where *ɛ* = 10^−8^ prevents division by zero.
8. **Peak voltage**: max(*b*).
9. **Trough voltage**: min(*b*).
10. **Arc-length** (line-length): 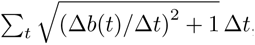, a waveform complexity measure sensitive to both amplitude and frequency [14].

Burst segments with duration *<* 10 ms or with zero apparent frequency (i.e. no detected peaks or troughs) are discarded, as they are too brief to yield reliable spectral features. All 10 features are then z-score normalised across all bursts of a given animal before clustering.

*c. Stage 3: Unsupervised clustering and automatic k selection.* The normalised feature matrix is clustered using *k*-means [17] with *k*-means++ initialisation. To determine the appropriate number of clusters *k* without manual inspection, we ran *k*-means for *k* = 2*, . . . ,* 19 and computed the normalised inertia (within-cluster sum of squared distances divided by the number of points) at each *k*. The optimal *k* was identified as the point of maximum curvature of the inertia curve, computed analytically as:

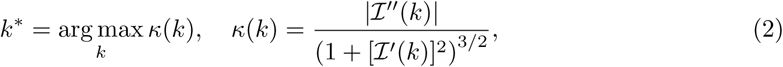

where I(*k*) is the normalised inertia and primes denote numerical derivatives. This maximum-curvature (elbow) criterion is fully automatic and gave *k* ∈ {6, 7, 8, 9} across the six animals (Table I).

Figure 1C shows a UMAP [21] projection of the 10-dimensional feature space for 30,000 randomly sampled bursts from FVB18. The eight cluster states occupy largely non-overlapping regions of the low-dimensional manifold, confirming that the *k*-means partition recovers morphologically coherent burst populations. The ribbon-like topology is consistent with the feature space lying near a low-dimensional manifold, principally parameterised by oscillation frequency (*r* = 0.51), with signal amplitude (std; *r* = 0.19) as a secondary contributor (data not shown), embedded in the 10-dimensional feature space. Figure 1D shows the burst duration distribution per cluster as violin plots, illustrating that states differ systematically in this single feature alone, with some states exclusively containing short bursts (*<* 50 ms) and others containing long, sustained events: state S6 spans a broad range extending up to ∼ 280 ms, while state S7 is concentrated at even longer durations, extending beyond 250 ms.

*d. Stage 4: State-vector construction.* Once cluster labels are assigned to every burst segment, the continuous LFP time series is converted to a discrete symbolic sequence. Each sample *t* within a seizure trial receives the label *s*(*t*) ∈ {−1, 0, 1*, . . . , k* − 1}: background samples retain the value −1, and each burst sample receives the cluster label of the burst it belongs to. This produces a trials × times integer matrix, the *state-vector array*, in which each row is a symbolic seizure trajectory. Figure 1B shows one complete seizure from FVB18, with the raw LFP overlaid with burst segments coloured by state. The shift in dominant colour from the early ictal period (states 2 and 4, cyan/green) to the late ictal period (states 0, 1, and 7, red/orange) is immediately apparent and motivates the Markov and entropy analyses that follow.

**Table 1.**
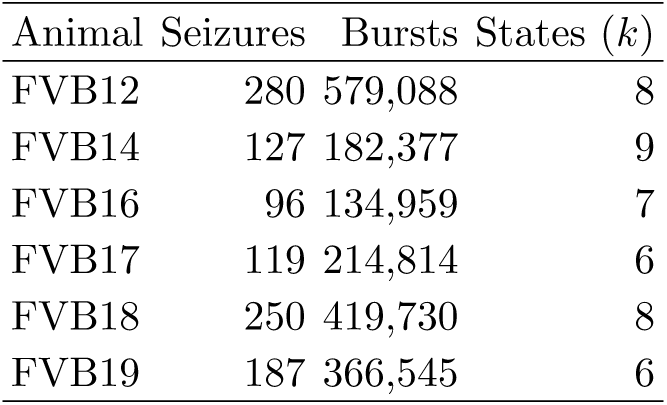
Summary of dataset and clustering results per animal. States (*k*) indicates the number of burst clusters selected by the elbow method.

| Animal | Seizures | Bursts | States ($k$) |
| --- | --- | --- | --- |
| FVB12 | 280 | 579,088 | 8 |
| FVB14 | 127 | 182,377 | 9 |
| FVB16 | 96 | 134,959 | 7 |
| FVB17 | 119 | 214,814 | 6 |
| FVB18 | 250 | 419,730 | 8 |
| FVB19 | 187 | 366,545 | 6 |

The full burst-detection and clustering pipeline is illustrated in Figure 1.

### 8. First-order Markov transition matrices

From the symbolic sequence *s*(*t*) we estimated the first-order Markov transition matrix **M** for each animal. The entry *M_ij_* is the empirical probability of transitioning from state *i* to state *j* in consecutive bursts:

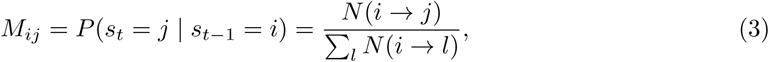

where *N* (*i* → *j*) is the count of all consecutive pairs (*i, j*) across all seizures. Self-transitions (*i* = *j*) were excluded by setting *M_ii_* = 0 before row normalisation, since within our burst-detection scheme two consecutive time points cannot carry the same label (bursts are contiguous regions, so a burst-to-itself transition is never recorded). All seizures of a given animal contributed to a single estimate of **M**.

### 9. Temporal evolution: Frobenius distance between consecutive epoch matrices

To quantify how rapidly the transition structure changes over the recording, we partitioned each animal’s seizure sequence into non-overlapping windows of *W* = 20 consecutive seizures and estimated a separate Markov matrix **M**^(*w*)^ for each window *w*. We then computed the Frobenius norm of the difference between adjacent matrices:

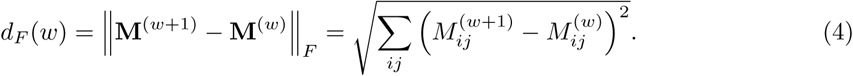

Large values of *d_F_* indicate that the transition structure underwent a large shift between two consecutive recording epochs. Each epoch was assigned a mean UTC hour from the seizure onset timestamps, enabling colour-coded visualisation of circadian phase.

This non-overlapping design discards information at window boundaries and yields coarse temporal resolution. We therefore adopted a sliding-window design as the primary analysis: the *W* = 20-seizure window is advanced by a stride of 1 seizure, and *d_F_* (*w*) is computed between each pair of adjacent sliding windows (which differ by exactly one seizure). This yields a substantially finer-grained trajectory (76–260 points per animal). Because consecutive sliding windows share 19 of 20 seizures, *d_F_* (*w*) values are strongly autocorrelated, so treating them as independent observations (as the non-overlapping design permits) would be anti-conservative. We therefore tested the group-level trend with a generalised estimating equations (GEE) model, *d_F_* ∼ window norm, with animal as the clustering variable and an AR(1) working correlation structure to account for this induced autocorrelation, in place of the linear mixed-effects model used for the non-overlapping design. As a supplementary check, we also computed *d_F_* for the disjoint (non-overlapping) windows directly from the sliding-window matrices (restricting to windows starting at multiples of 20 seizures); this gives estimates numerically identical to the disjoint-window analysis reported alongside the sliding-window results (Results; Figure 5), confirming that the two computational routes are consistent.

A state with few observed outgoing transitions within a given window yields a highly unstable row of **M**^(*w*)^: with zero observations the row is undefined (set to all zero), and with a single observation it is a one-hot vector regardless of the state’s true transition preferences. As such a row enters or leaves the window, it can toggle between these extremes and contribute up to its own norm (∼ 1, since rows are probability vectors) to *d_F_* , a sampling artefact of the window’s finite seizure count rather than a genuine change in transition structure. We therefore masked each row of **M**^(*w*)^ from the *d_F_* comparison unless the corresponding state had at least 10 observed outgoing transitions in *both* windows being compared. This threshold was set from the empirical distribution of per-row observation counts (median 1,574 observations per row-window; 11.7% of rows have zero observations and a further 5.5% fall between 1 and 9, with a natural gap separating this sparse tail from the well-sampled majority) before inspecting any resulting trend statistic, and the group-level GEE result below is stable in sign and significance for thresholds between 3 and 30 observations.

### 10. Sequential KL divergence between adjacent seizures

To characterise how similar the burst-type occupancy distribution of each seizure is to that of the preceding seizure, we computed the symmetric Kullback-Leibler (KL) divergence between consecutive seizure state histograms. For seizures *i* − 1 and *i* with empirical state distributions *p* and *q*:

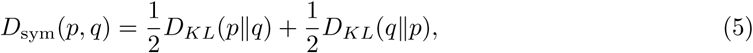

with 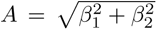and a small additive constant *ε* = 10^−10^ to handle empty bins. Consecutive values were pooled into *W* = 20-seizure bins; the bin mean and standard error of the mean are shown. A three-bin moving average is overlaid to highlight trends. Following the same sliding-window rationale as the Frobenius analysis above, we replaced this disjoint binning with a rolling mean and SEM (window = 20 seizures, stride = 1, centred, minimum 1 observation) as the primary display, since the underlying KL divergence is already computed at full seizure-pair resolution and disjoint pooling discards temporal information unnecessarily.

### 11. Burst-pattern entropy and circadian modulation

For each seizure, we computed the normalised Shannon entropy of the burst-type distribution:

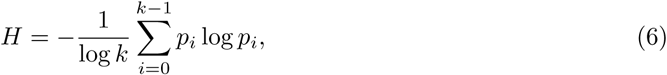

where *p_i_* is the fraction of detected bursts in seizure *s* belonging to state *i*, and *k* is the number of states. By dividing by log *k*, *H* ranges from 0 (all bursts in one state) to 1 (uniform distribution over states). Each seizure was assigned to one of five time-of-day periods based on the UTC hour of its onset: Night (22:00–05:59), Early Morning (06:00–09:59), Morning (10:00–13:59), Afternoon (14:00–17:59), and Late Afternoon (18:00–21:59). Differences in *H* across periods were tested with the Kruskal-Wallis test; pairwise comparisons used the Mann-Whitney U test with Holm-Bonferroni correction for multiple comparisons.

Because the five-period binning discretises a continuous circadian phase, we additionally fitted a single-component 24-hour cosinor model [23] to the per-seizure entropy of each animal: *H*(*t*) = *M* + *β*_1_ cos(2*πt/*24) + *β*_2_ sin(2*πt/*24) + *ε*, where *t* is the fractional UTC onset hour. The amplitude of circadian modulation is *A* = ^√^*β*^2^ + *β*^2^ and the acrophase (peak time) is *φ* = atan2(*β*_2_*, β*_1_), converted to a peak hour as *φ* · 24*/*(2*π*) mod 24. Significance of the circadian component was assessed with the joint F-test of *β*_1_ = *β*_2_ = 0 (the standard cosinor zero-amplitude test), fitted by ordinary least squares per animal.

We separately characterised the circadian distribution of seizure occurrence itself (as opposed to within-seizure entropy) using 24-hour polar histograms of onset hour per animal (Supplementary Figure S2), and tested each animal’s onset-hour distribution for circular non-uniformity with the Rayleigh test [15], which returns the mean resultant length *R* (0 = uniform, 1 = all onsets at a single hour) and an associated *p*-value.

### 12. Higher-order Markov analysis: block-entropy rate

To test whether burst sequences contain memory beyond the first-order Markov assumption, we computed the block-entropy rate increment *F_N_* as a function of history length *N* [9]:

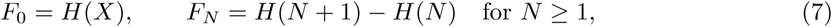

where *H*(*N* ) denotes the Shannon entropy (in bits) of all length-*N* n-grams in the burst sequence. For a purely first-order Markov process, *F_N_* is constant for all *N >* 1. A monotonically decreasing *F_N_* indicates that longer burst histories provide additional predictive power, signalling the presence of higher-order dependencies. We computed *F_N_* for *N* = 0*, . . . ,* 5 on the full burst sequence of each animal, and separately on the early, middle, and late thirds of each animal’s seizure record (defined by seizure index).

### 13. Statistical analysis

Group differences in entropy across time-of-day periods were assessed with the Kruskal-Wallis H test (non-parametric one-way ANOVA). Pairwise post-hoc tests used the Mann-Whitney U test with Holm-Bonferroni family-wise error correction. Group-level trends in the sliding-window Frobenius distance were assessed with a generalised estimating equations (GEE) model with an animal-clustered AR(1) working correlation structure (above); circadian modulation of entropy was additionally assessed by cosinor regression, and circadian clustering of seizure occurrence by the Rayleigh test (above). All analyses were performed in Python using numpy, scipy, pandas, statsmodels, seaborn, and xarray.

### 14. Animal classification from burst-type usage

To test whether the pooled burst-type vocabulary carries an animal-identity signature, we used the same *N* = 1,897,513 pooled, standardised burst feature vectors as in the common-space analysis (Supplementary Figure S3) and re-clustered them with MiniBatchKMeans (mini-batch size 10,000, 10 restarts, fixed random seed for reproducibility), again selecting the number of burst types by the same maximum-curvature (elbow) criterion described above, which yielded *n* = 6 types.

For each seizure we then built a burst-type usage profile – the normalised frequency of each of the six burst types among that seizure’s bursts – and trained a random-forest classifier (300 trees, balanced class weights) to predict the source animal from this profile alone. Out-of-fold predictions were obtained via stratified 5-fold cross-validation (fold assignment fixed by seed), and classifier performance was summarised as balanced accuracy against chance level (1*/*6). Feature importances were averaged across folds to rank burst types by their contribution to animal discrimination, and the top three most discriminative types were further characterised by (i) their timing relative to seizure onset (kernel density estimate of burst onset time within the peri-ictal window) and (ii) their circadian distribution (time-of-day histogram of burst occurrence).

## RESULTS

### 15. Seizures are organised by recurring burst-state sequences

A central unresolved question in epilepsy is how seizure dynamics evolve over the course of disease progression. Specifically, it remains unclear whether pathological electrophysiological bursts follow reproducible trajectories from one seizure to the next, and how such temporal patterns emerge during epileptogenesis and become established in the chronic epileptic state on a subject-specific basis. To address this, we developed a data-driven pipeline (Figure 1; Methods) that converts continuous LFP recordings from six pilocarpine-treated FVB mice into discrete symbolic sequences of burst types. Adaptive *z*-score segmentation, followed by 10-dimensional morphological feature extraction and unsupervised *k*-means clustering with automatic model-order selection, identified between *k* = 6 and *k* = 9 recurrent burst states per animal (Table I), yielding 134,959-579,088 labelled burst events per animal, a corpus large enough to support robust statistical inference at every stage of the analysis.

Visualising the resulting sequences as state-vector matrices (Figure 2), in which each row is a single seizure colour-coded by burst type and rows are stacked chronologically, immediately establishes that intra-seizure dynamics are far from random. Every animal exhibits rich burst-type diversity within individual episodes: the LFP consistently traverses a substantial fraction of the available state alphabet during each ictal event, rather than staying confined in a single dominant discharge pattern. More importantly, this diversity is temporally organised. Certain burst types predominate in the early ictal period, often within seconds of onset, while others emerge preferentially during the sustained discharge or at offset, producing horizontal colour bands that recur with high fidelity across seizures (vertical coherence in Figure 2). This sequential structure is especially pronounced in FVB16 and FVB19, where a sharp compositional transition in dominant burst type occurs approximately 20–40 s after onset, suggestive of a reproducible two-phase ictal architecture. At the same time, the specific colour palettes and temporal arrangements differ markedly between animals, reflecting individual differences in the burst morphologies and network architectures shaped by each animal’s epileptogenic process. These observations establish the foundational result of this study: seizures in the pilocarpine FVB model are not monolithic events but are composed of structured, reproducible sequences of morphologically distinct burst types whose arrangement encodes information about the state of the epileptic network.

**Figure 2.**
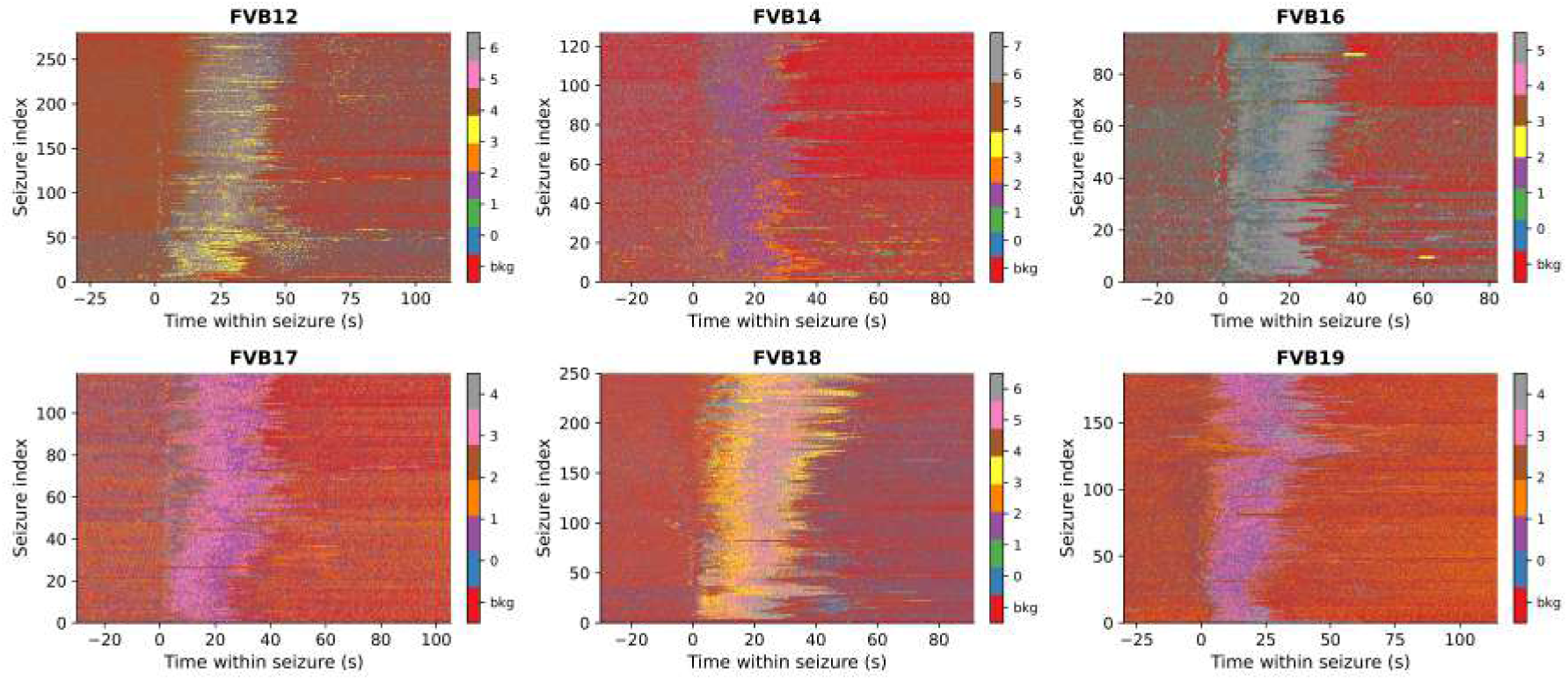
Seizure state vectors reveal rich, temporally structured burst-type sequences. Each panel shows one animal (FVB12–FVB19). Rows are individual seizures ordered chronologically from bottom to top; columns are time within the seizure (s) aligned to onset (0 s, vertical boundary between pre-ictal and ictal activity). Colour encodes burst type (0 to *k* − 1, see colourbar); red indicates background (sub-threshold) activity with no detected burst (label −1). The pre-onset epoch (−25 to 0 s) is predominantly red, confirming that burst detection correctly captures the onset boundary. After onset, multiple distinct burst types are recruited within every seizure, with visible temporal structure: certain states predominate early in the ictal period while others emerge later, producing horizontal colour bands that indicate sequential dynamics rather than random sampling. This organisation is especially prominent in FVB16 and FVB19, where a shift in dominant colour occurs 20–40 s after onset. Across animals, the same broad structure recurs seizure-to-seizure (vertically coherent colour patterns), yet the specific palette and its temporal arrangement differ markedly between animals, reflecting animal-specific burst morphologies shaped by each individual’s epileptogenic network. The total number of seizures and burst-state count *k* per animal are given in Table I.

Quantification of burst-state composition across the three peri-ictal phases confirms this picture at the population level (Supplementary Figure S1): background activity dominates the pre-ictal window in all animals, burst-state occupancy rises sharply at onset, and the ictal-to-post-ictal transition involves a partial, but incomplete and animal-specific, return toward the pre-ictal composition.

The observation of structured burst-type sequences raises an immediate mechanistic question: are these sequences governed by consistent statistical rules, or do they merely reflect a broad tendency for certain morphologies to dominate at different phases of a seizure without constraining the specific burst-to-burst transitions? To answer this, we estimated first-order Markov transition matrices from the full seizure records of each animal, pooling across all episodes to obtain stable empirical estimates of the probability *M_ij_* that burst type *j* follows immediately after burst type *i* (Equation 3; Figure 3).

**Figure 3.**
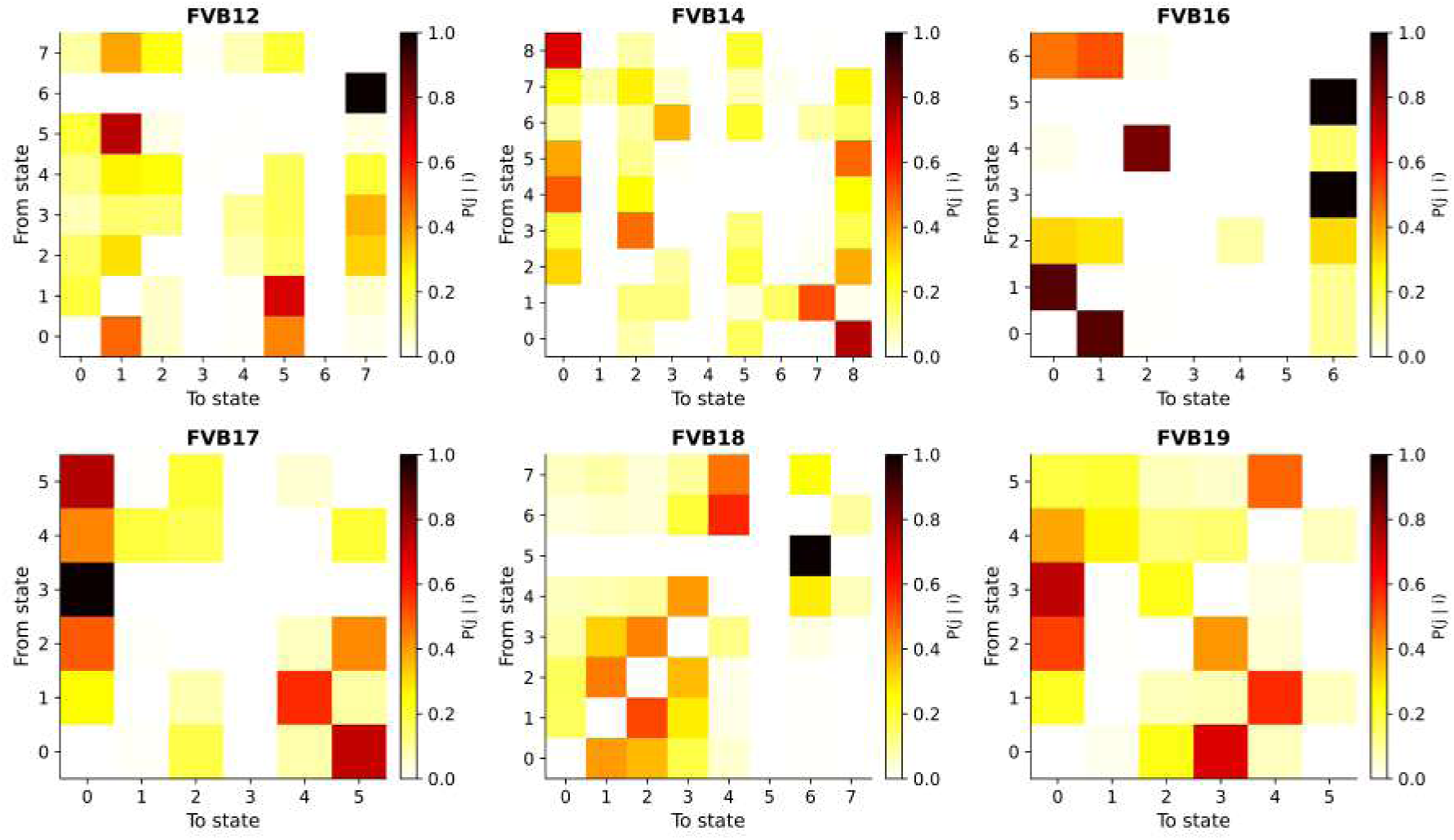
First-order Markov transition matrices reveal sparse, animal-specific burst grammars. Each panel shows the row-normalised transition probability matrix **M** estimated from all seizures of one animal. Entry *M_ij_* is the empirical probability of observing burst type *j* immediately after burst type *i*. Self-transitions are excluded (diagonal forced to zero). The shared colourscale runs from 0 (white, no observed transition) to 1.0 (black, deterministic transition); the same scale is used across all animals to enable direct comparison. Despite the large number of observed bursts (1.3 × 10^5^ to 5.8 × 10^5^ per animal; Table I), the matrices are markedly sparse: the majority of off-diagonal entries remain near zero, indicating that each burst type transitions predominantly to a small subset of successor states. The dominant targets differ by animal, in FVB12 states 1 and 5 accumulate high probability mass from several source states, with state 7 also receiving strong convergent input; in FVB16 states 0, 1, and 6 act as strong attractors with transition probabilities exceeding 0.8; FVB14 shows state 0 and state 8 as preferred sinks. FVB18 is comparatively more distributed, with moderate probabilities spread across several pairs. This inter-animal heterogeneity in transition topology, computed independently for each animal, reflects genuine differences in the network architecture established during the epileptogenic process rather than a measurement artefact.

The result is unambiguous: the transition matrices are markedly sparse. Despite very large numbers of observed burst-to-burst transitions (up to 5.8 × 10^5^ per animal), the vast majority of off-diagonal entries remain close to zero. Each burst type transitions predominantly to only one to three successor states with substantial probability, while transitions to all other states are essentially absent. This sparsity is not a consequence of limited sampling – these are richly sampled datasets – but instead reflects genuine constraints on sequential dynamics: the epileptic network does not sample randomly from its state alphabet but follows a constrained set of preferred pathways, a structured grammar of burst-type successions [6**?** ].

The topology of this grammar is animal-specific in a reproducible way. In FVB12, states 1 and 5 function as strong attractors, drawing high-probability transitions from multiple source states, and state 7 additionally receives strong convergent input, including a near-deterministic transition from state 6. In FVB16, states 0, 1, and 6 dominate as near-deterministic hubs with transition probabilities exceeding 0.8. FVB14 channels most transitions toward state 0 and state 8 as preferred sinks, while FVB18 shows a more distributed pattern with moderate probabilities spread across several state pairs. Because burst morphologies and cluster labels were derived entirely independently for each animal, these inter-individual differences in transition topology cannot be attributed to representational artefact; they reflect genuine variability in the biological substrate, such as the synaptic weights or excitability profiles, established during each animal’s epileptogenic process.

### 16. Seizure grammar stabilises with chronicity

Having established that burst-type transitions are constrained at any given moment, we asked whether this constraint itself evolves over the weeks-long chronic recording, whether the grammar is stationary or whether it undergoes systematic reorganisation as the epileptic network matures. To capture this, we partitioned each animal’s seizure sequence into non-overlapping windows of *W* = 20 consecutive seizures and tracked the Frobenius norm of the difference between adjacent transition matrices, *d_F_* (*w*) (Equation 4), a continuous measure of how rapidly the grammar is changing at each point in the recording.

Using the sliding-window design with under-sampled transition-matrix rows masked (Methods), the Frobenius distance trajectory was derived in a finer temporal grain series (Figure 4). Across all six animals, the smoothed (9-window moving-average) trajectory trends downward, and Kendall’s *τ* is consistently negative, though modest in magnitude (FVB12: *τ* = −0.13; FVB14: *τ* = −0.12; FVB16: *τ* = −0.36; FVB17: *τ* = −0.21; FVB18: *τ* = −0.12; FVB19: *τ* = −0.24) – a more internally consistent picture across animals than the noisier disjoint-window design produced. We note that specific single-window outliers highlighted by the coarse disjoint-window design are not a reliable feature of the data: in FVB14, for example, the largest *d_F_* occurs at the fifth (final) disjoint window rather than the first, and no single window’s distance is more than double any other (Figure 5) – underscoring why we treat the finer-grained, appropriately corrected sliding-window analysis as primary when drawing conclusions about any individual animal’s trajectory, with the disjoint-window design retained in Figure 5 as a supplementary comparison.

**Figure 4.**
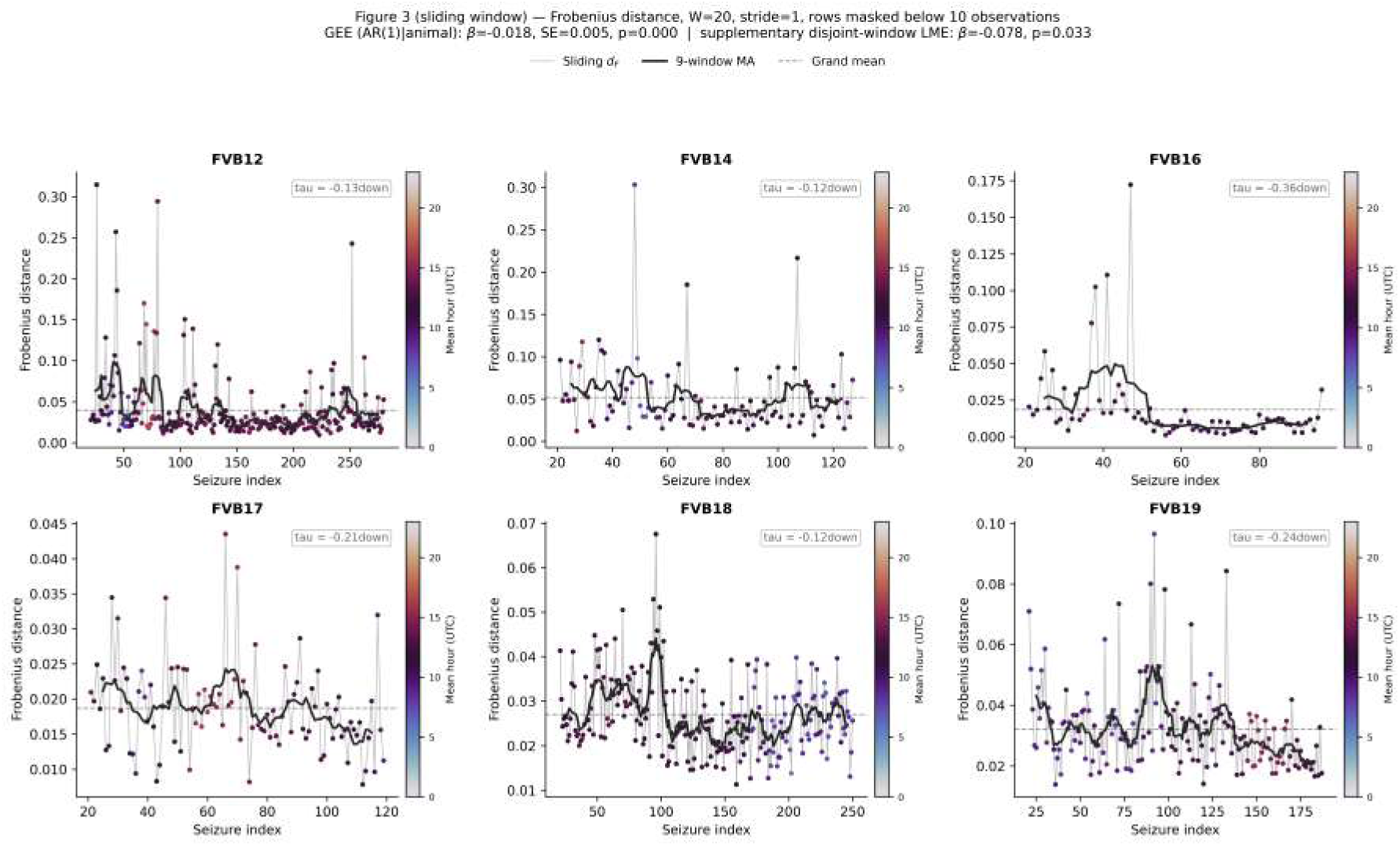
Temporal profile of Markov transition-matrix change across the chronic recording (sliding-window design). Each point shows the Frobenius distance *d_F_* (*w*) between the first-order Markov matrices of two adjacent sliding windows of *W* = 20 seizures (stride = 1 seizure; rows with fewer than 10 observed transitions in either window masked from the comparison, Methods), plotted at the seizure index of the end of the right-hand window. Points are coloured by the mean UTC onset hour of the right-hand window (twilight colourmap: dark purple = night, light yellow = midday). The thin grey line connects consecutive measurements; the black curve is a 9-window moving average; the dashed horizontal line marks each animal’s grand mean *d_F_* . The inset in each panel shows Kendall’s *τ* between window order and *d_F_* as a descriptive measure of trend direction; no per-animal inference is intended. A GEE model (animal-clustered, AR(1) working correlation, accounting for the autocorrelation induced by window overlap; Methods) pooling all animals yielded *β*^^^ = −0.018 (SE = 0.005, *p* = 1.3 × 10^−4^, *n* = 939 windows): a significant declining group-level trend. Individual *τ* values are uniformly negative but modest (FVB12: −0.13; FVB14: −0.12; FVB16: −0.36; FVB17: −0.21; FVB18: −0.12; FVB19: −0.24), a more internally consistent picture than the disjoint-window design produced (Figure 5).

**Figure 5.**
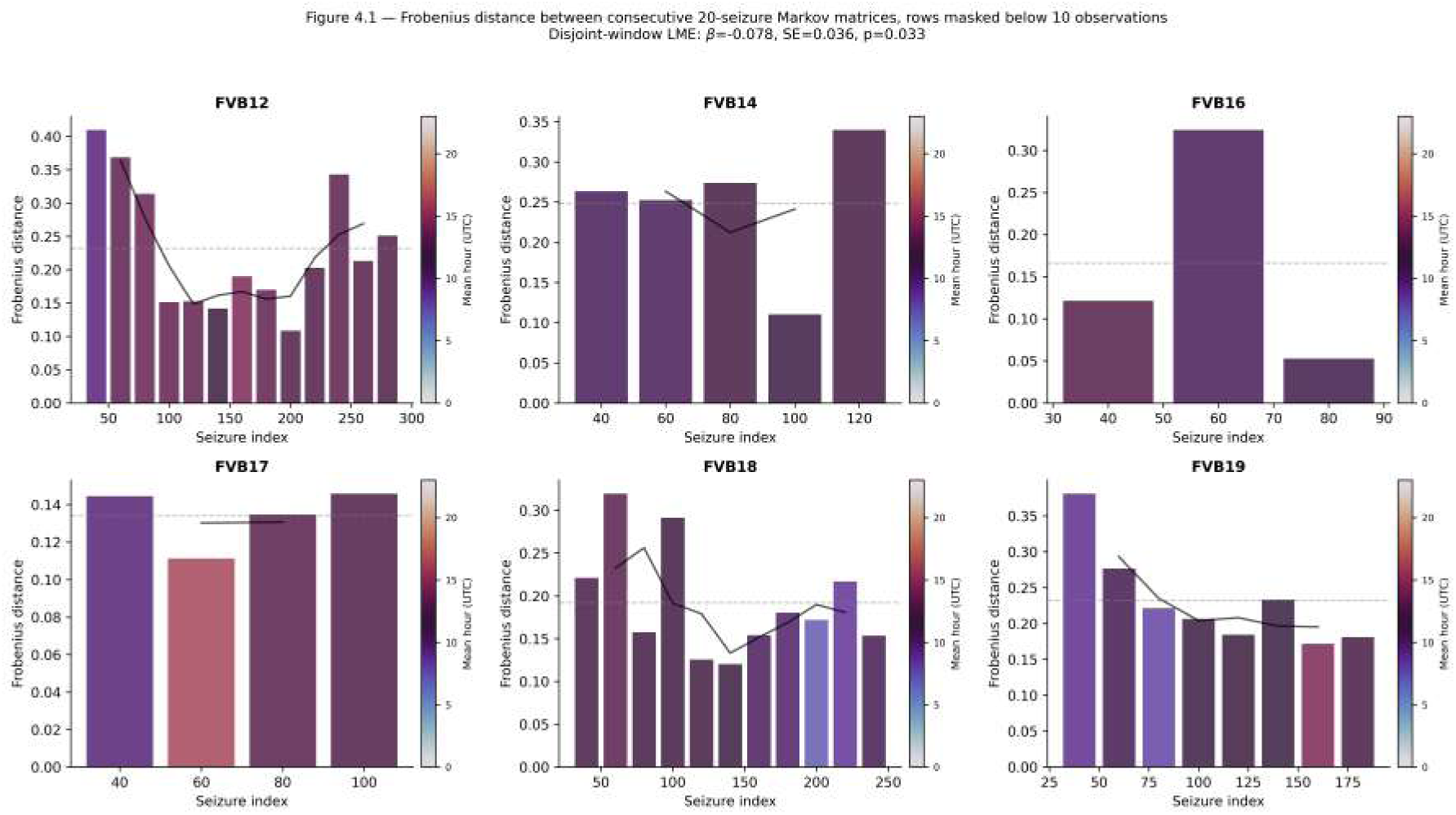
Frobenius distance between consecutive non-overlapping 20-seizure Markov matrices (disjoint-window comparison). Same underlying (row-count-masked) transition matrices as Figure 4, but restricted to genuinely disjoint (non-overlapping) 20-seizure windows, presented as a supplementary comparison to the sliding-window design (Methods). Bar height is *d_F_* (*w*) for each consecutive pair of disjoint windows; bars are coloured by mean UTC onset hour of the right-hand window (twilight colourmap). The black curve shows a 3-window moving average. A linear mixed-effects model on these disjoint windows gives *β*^^^ = −0.078 (SE= 0.035, *p* = 0.026, *n* = 44; estimate stable across optimisers, *p* = 0.02–0.03); this estimate is numerically identical whether *d_F_* is computed directly on disjoint windows or extracted from the sliding-window matrices used in Figure 4. Note that the tallest bar is not always the first: in FVB14, for example, the fifth (final) window carries the largest distance, not the first, and no single window is more than double any other – the apparent ”early peak” visible by eye in this coarse, low-resolution design is sensitive to which few windows happen to be sampled, which is why the sliding-window analysis in Figure 4 is treated as the primary result.

Because consecutive sliding windows share 19 of 20 seizures, *d_F_* (*w*) is strongly autocorrelated; we therefore tested the group-level trend with a GEE model (animal-clustered, AR(1) working correlation; Methods) rather than treating sliding-window points as independent observations. With under-sampled rows masked (Methods), this yielded *β*^^^ = −0.018 (SE = 0.005, *p* = 1.3 × 10^−4^, *n* = 939 sliding windows, AR(1) *α* = 0.36): a significant decline in transition-matrix change across the chronic recording. As a supplementary comparison, applying the coarser disjoint (non-overlapping) 20-seizure windowing scheme to the same masked transition matrices and fitting a linear mixed-effects model gives *β*^^^ = −0.078 (SE = 0.035, *p* = 0.026, *n* = 44; Figure 5; point estimate stable across optimisers, *p* = 0.02–0.03). The two designs agree in direction and significance: the sliding-window/GEE combination provides a more appropriately powered and less noise-sensitive test of the same question, with a correspondingly smaller point estimate. Between-animal heterogeneity remains substantial regardless of windowing scheme (animals differ in the absolute level of *d_F_* , from a mean of 0.019 in FVB16 to 0.052 in FVB14, and in the shape of their temporal trajectories). Colour-coding data points by time of day reveals no consistent circadian patterning in the Frobenius distances, arguing against diurnal excitability fluctuations as the primary driver.

Taken together, these data indicate a modest but statistically robust, animal-general decline in transition-matrix change across the chronic recording, significant at the group level under both the primary sliding-window/GEE test and the supplementary disjoint-window LME. We remain cautious, however, about attributing this pattern to a specific stage-limited process, such as a discrete early phase of heightened network reconfiguration, as opposed to a slow, continuous drift: the effect size is small relative to the between-animal and within-animal variability visible in Figure 4, and the six per-animal *τ* values, while uniformly negative, span a threefold range (−0.12 to −0.36). Either interpretation is consistent with ongoing network reorganisation during epileptogenesis (e.g. synaptic remodelling, mossy fibre sprouting, or inhibitory interneuron loss [8, 12, 13, 25]); distinguishing a discrete reconfiguration phase from continuous drift would require a larger cohort with standardised recording durations and denser early-phase sampling than the present six animals provide.

### 17. Consecutive seizures diverge maximally in burst-type content early in the chronic phase, then converge

The Frobenius analysis captures changes in transition *topology* ; it does not directly address whether the dominant burst types active within individual seizures also shift over time. To probe this complementary dimension of non-stationarity, we computed the symmetric Kullback-Leibler (KL) divergence between the burst-type occupancy distributions of every pair of consecutive seizures (Equation 5) and pooled values into 20-seizure bins. This measure asks a simpler but equally fundamental question: are two back-to-back seizures compositionally similar, visiting the same burst-state repertoire in roughly the same proportions?

The answer depends strongly on when in the recording those seizures occur (Figure 6, shown here as a stride-1 rolling mean over the same *W* = 20-seizure window used for the Frobenius analysis, rather than disjoint bins – Methods). KL divergence is highest within the first ∼ 20 seizures in most animals, often by a large margin: FVB12 reaches ≈ 0.74 nats early and *<* 0.15 nats thereafter, while FVB14 declines from ≈ 0.44 nats to ≈ 0.10 nats. Early in the chronic phase, therefore, consecutive seizures can be compositionally radically different, one episode dominated by high-frequency, short-duration states while the next is dominated by sustained low-frequency discharge, reflecting a network that has not yet settled into a preferred ictal mode. As the recording progresses, consecutive seizures converge on more similar occupancy distributions, pointing toward a stereotyped compositional signature.

**Figure 6.**
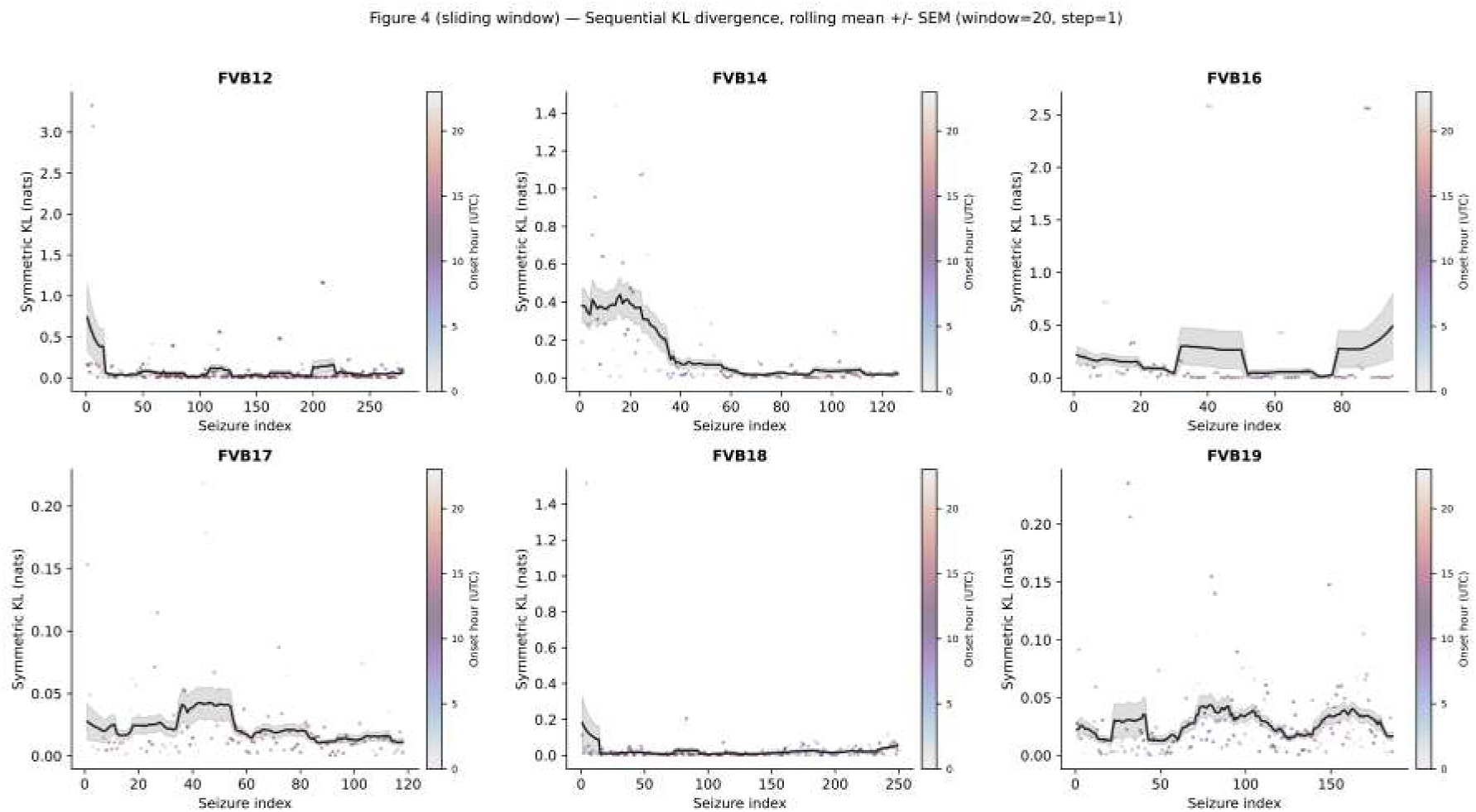
Sequential KL divergence between consecutive seizures reveals an early transient of maximal state-space dissimilarity that decays as epilepsy matures (rolling-window design). Symmetric KL divergence (nats; Equation 5) between burst-type occupancy distributions of consecutive seizure pairs (small points), with a rolling mean ±SEM (window = 20 seizures, stride = 1; Methods) overlaid as a black line with shaded band. Colour: onset hour of the seizure pair. The rolling mean peaks within the first ∼ 20 seizures in most animals (FVB12: ≈ 0.74 nats; FVB14: ≈ 0.44 nats), indicating that early consecutive seizures sample markedly different burst-type repertoires. Divergence declines with disease progression, consistent with convergence toward a stereotyped compositional signature. FVB17, FVB18, and FVB19 show comparatively low divergence throughout (*<* 0.05 nats for FVB17 and FVB19; a brief early transient up to ≈ 0.18 nats in FVB18 before settling to a low plateau). FVB19 exhibits a mild secondary rise (to ≈ 0.04 nats) near seizure 75–80, indicating transient renewed variability.

Three animals (FVB17, FVB18, FVB19) show comparatively low KL divergence throughout (*<* 0.05 nats for FVB17 and FVB19, aside from a brief early transient up to ≈ 0.18 nats in FVB18), suggesting that their networks either stabilised rapidly after SE or that recording began only during the consolidated phase. FVB19 provides a subtle exception: a mild secondary rise near seizure 75–80 suggests a transient episode of renewed compositional variability before eventual restabilisation.

The convergence between the Frobenius and KL results is notable because the two metrics are conceptually orthogonal: one probes the rules by which burst states succeed one another, the other probes which burst states are present – yet they tell a consistent story. Early in the chronic phase, the seizure grammar is unstable at multiple levels of description simultaneously; this instability declines as the disease progresses, suggesting that the epileptic network consolidates toward a deeper, more stereotyped attractor.

### 18. The circadian cycle shapes the internal dynamics of individual seizures in a subset of animals

Circadian rhythms are well-established modulators of seizure timing: the probability of seizure occurrence fluctuates reliably with time of day across species and models [1, 20]. Whether circadian phase also shapes the internal dynamics of individual ictal episodes – whether a seizure occurring at night is dynamically distinct from one occurring at noon – is a far less explored question. To address it, we computed the normalised Shannon entropy *H* of the burst-type occupancy distribution within each seizure (Equation 6), a single-number index of within-episode burst-type diversity (*H* = 0: all bursts of one type; *H* = 1: perfectly uniform across all *k* states), and tested whether *H* varied systematically across five time-of-day periods.

Two of six animals show statistically significant circadian modulation of within-seizure entropy (Kruskal-Wallis: FVB12, *p* = 0.008; FVB14, *p* = 0.008; Figure 7). In FVB12, Night-time seizures carry significantly higher entropy than Morning seizures (Holm-Bonferroni-corrected Mann-Whitney, *p <* 0.01): nocturnal episodes recruit a broader, less stereotyped repertoire of burst types. In FVB14, the effect is directionally reversed: Night-time seizures have significantly lower entropy than Late Afternoon seizures (Holm-corrected *p <* 0.05), indicating that nocturnal episodes in this animal are the more stereotyped, not the more diverse.

**Figure 7.**
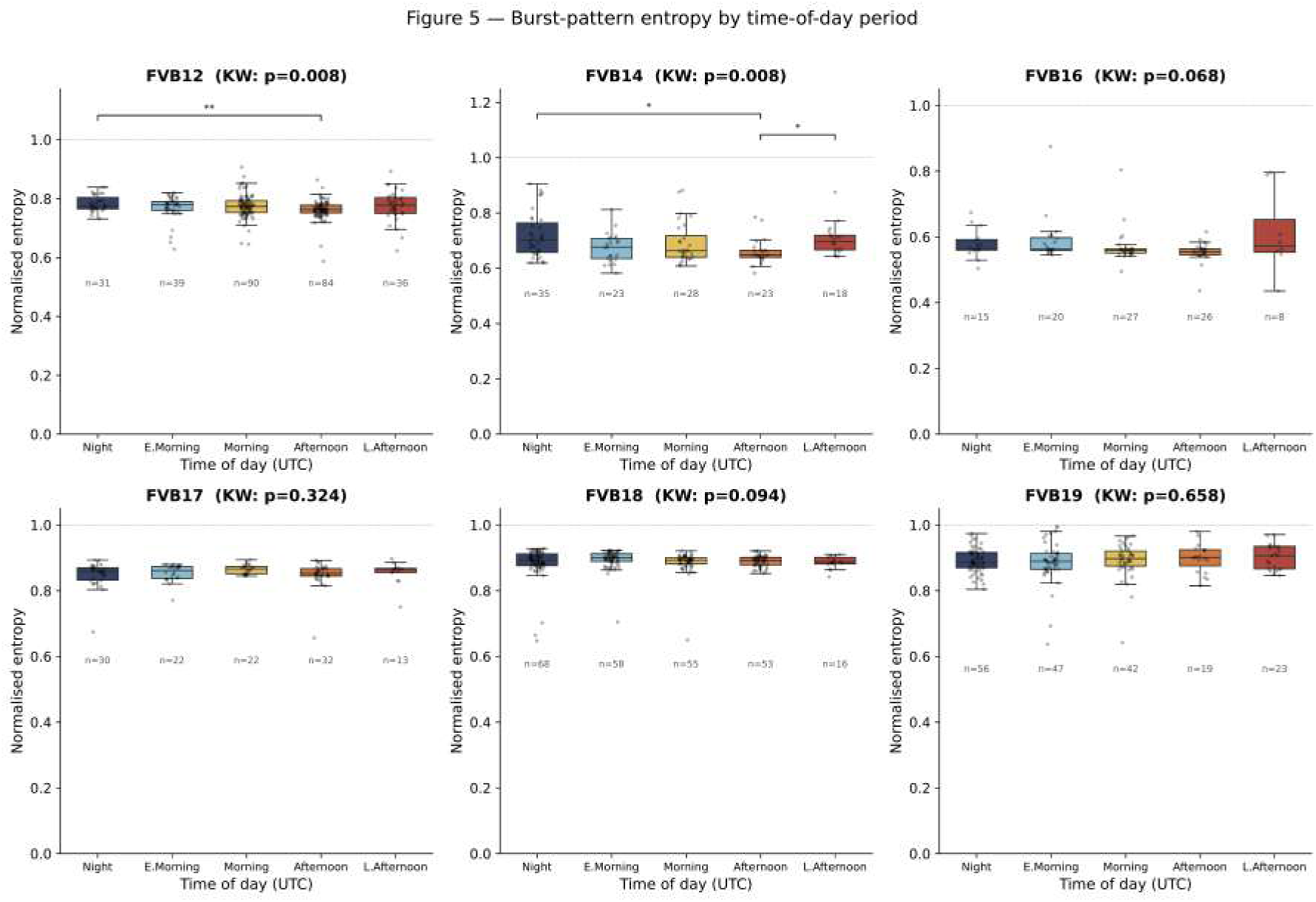
Within-seizure burst-pattern entropy is modulated by circadian phase in two of six animals. Normalised Shannon entropy *H* (Equation 6) of the burst-type occupancy distribution per seizure, grouped into five UTC time-of-day bins: Night (22:00–05:59), Early Morning (06:00–09:59), Morning (10:00–13:59), Afternoon (14:00–17:59), Late Afternoon (18:00–21:59). Box plots: median and IQR; strip plots: individual values; *n* per bin printed below each box. Subplot titles: Kruskal-Wallis omnibus *p*-value. Brackets: pairwise contrasts surviving Holm-Bonferroni correction (^∗^*p <* 0.05; ^∗∗^*p <* 0.01). FVB12 (*p* = 0.008): Night entropy significantly elevated relative to Morning, indicating more diverse nocturnal dynamics. FVB14 (*p* = 0.008): Night entropy significantly lower than Late Afternoon, indicating more stereotyped nocturnal dynamics. The opposing directions of modulation across animals indicate animal-specific circadian entrainment of within-seizure dynamics.

This animal-to-animal difference in the direction of circadian modulation is itself informative. It argues against a simple model in which circadian drive uniformly increases or decreases burst diversity, and instead points to an interaction between circadian phase and individual network architecture that depends on how each animal’s epileptogenic reorganisation has configured the excitability landscape.

The four animals that do not reach significance (FVB16, *p* = 0.068; FVB17, *p* = 0.324; FVB18, *p* = 0.094; FVB19, *p* = 0.658) include two (FVB16 and FVB18) that trend toward significance but are likely underpowered: the smallest time-of-day groups in these animals contain fewer than 20 seizures per period.

Because the five-bin Kruskal-Wallis test discretises a continuous circadian phase, we cross-checked this result with a cosinor regression (Methods; Figure 8) fitted directly to onset hour. The cosinor test agrees closely with the Kruskal-Wallis result – FVB12 (amplitude *A* = 0.010, *p* = 0.038) and FVB14 (*A* = 0.024, *p* = 0.021) again reach significance and no other animal does, with the exception of FVB19, which registers a near-miss under the cosinor test (*A* = 0.013, *p* = 0.058) that the coarser five-bin test did not detect at all (*p* = 0.658) – consistent with a smooth, graded circadian effect in FVB19 that a discrete binning can under-power. We take the convergence of two different circadian statistics on the same two animals as reassurance that the FVB12/FVB14 effect is not an artefact of the specific binning scheme.

**Figure 8.**
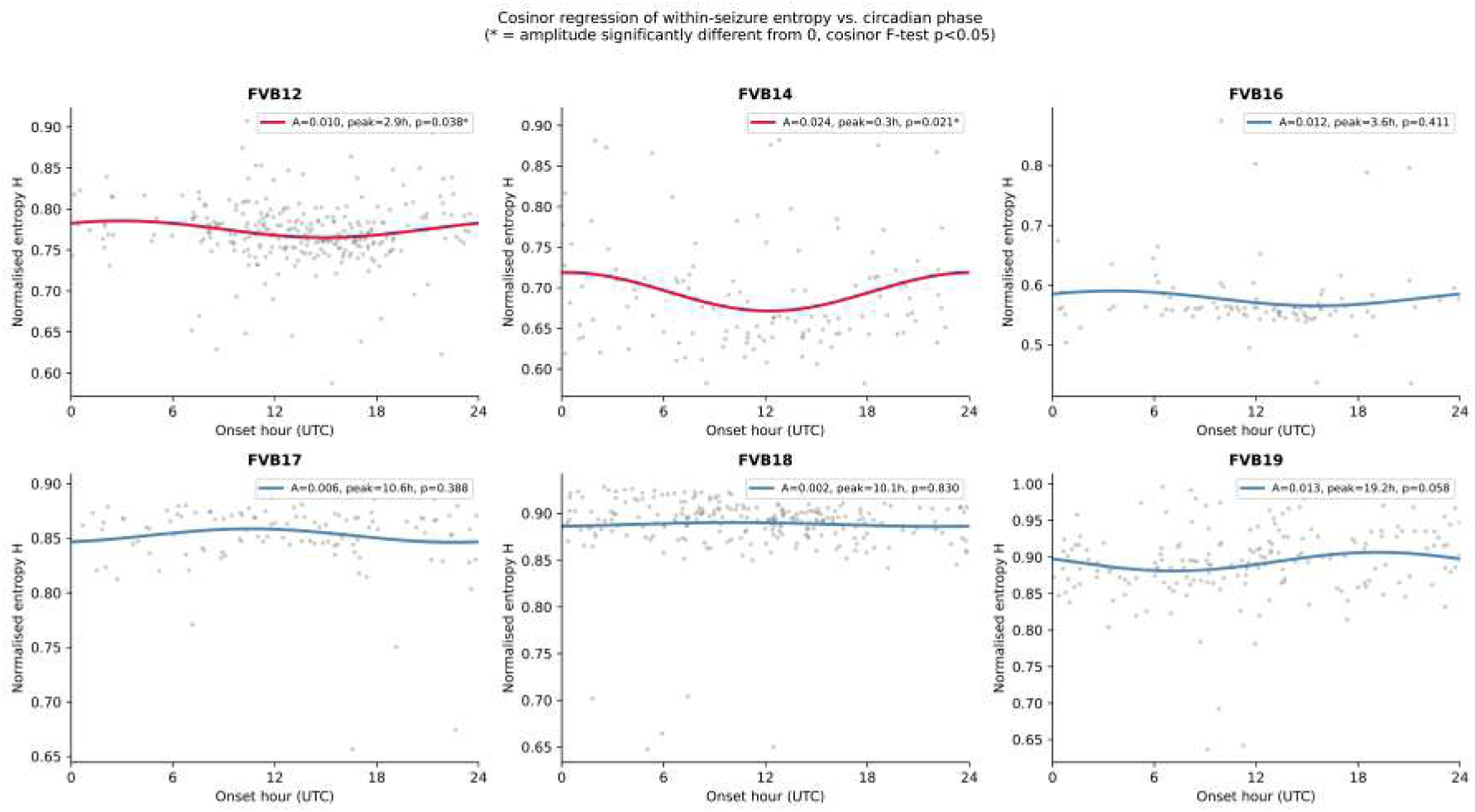
Cosinor regression of within-seizure entropy against circadian phase confirms the five-bin Kruskal-Wallis result. Per-seizure normalised entropy *H* (grey points) plotted against fractional UTC onset hour, with the fitted single-component 24-hour cosinor model overlaid (Methods): *H*(*t*) = *M* + *β*_1_ cos(2*πt/*24) + *β*_2_ sin(2*πt/*24). Curve coloured red where the amplitude is significantly different from zero (cosinor *F* -test, *p <* 0.05), blue otherwise. Legend reports amplitude 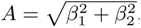, peak (acrophase) hour, and *p*-value per animal. FVB12 (*A* = 0.010, peak 2.9 h UTC, *p* = 0.038) and FVB14 (*A* = 0.024, peak 0.3 h UTC, *p* = 0.021) reach significance, matching the Kruskal-Wallis result in Figure 7. FVB19 is a near-miss (*A* = 0.013, *p* = 0.058) not detected by the coarser five-bin test.

We also directly tested the circadian distribution of seizure *occurrence itself* (as opposed to within-seizure entropy) using 24-hour polar histograms and the Rayleigh test for circular non-uniformity (Methods; Supplementary Figure S2). Seizure timing is significantly non-uniform across the day in four of six animals (FVB12, *p* = 1.1×10^−23^; FVB16, *p* = 3.2×10^−6^; FVB18, *p* = 7.6×10^−7^; FVB19, *p* = 0.044), but not in FVB14 (*p* = 0.124) or FVB17 (*p* = 0.081). Notably, this is a different set of animals from those showing significant circadian modulation of within-seizure entropy: FVB14 has strongly circadian entropy despite comparatively uniform seizure timing, whereas FVB16 and FVB18 show strongly non-uniform seizure timing without a corresponding entropy effect. This dissociation indicates that circadian gating of *when* seizures occur and circadian modulation of *how* a seizure unfolds internally are at least partially independent phenomena, each with its own animal-specific expression.

Taken together, these findings extend the circadian question from when seizures happen to how they unfold, establishing that the same biological clock mechanism that gates seizure probability also partially entrains the internal dynamical grammar of individual ictal episodes [20**?** ].

### 19. Burst sequences carry higher-order temporal memory whose depth and structure evolve with disease progression

The sparse first-order Markov structure establishes that burst-type transitions are constrained; but does the immediately preceding burst state fully capture that constraint, or do longer burst histories carry additional predictive power? This question is mechanistically important because genuine higher-order memory would imply that the ictal network operates with a form of temporal context, that the likelihood of the next burst state is determined not merely by the last burst but by the trajectory leading up to it, perhaps reflecting the accumulated effects of post-burst refractoriness, depletion of synaptic resources, or short-term excitability fluctuations. To test this, we computed the block-entropy rate increment *F_N_* as a function of history length *N* (Equation 7): for a strictly first-order Markov process *F_N_* is constant for *N* ≥ 1, whereas a monotonic decline with increasing *N* constitutes evidence of higher-order temporal memory.

The answer is unequivocal: in all six animals, *F_N_* declines monotonically from *N* = 0 to *N* = 5 (Figure 9). Burst sequences are not first-order Markov. Knowing the two or three burst types that preceded the current one reduces uncertainty about the next state beyond what the single immediately preceding burst alone provides, demonstrating that intra-seizure dynamics possess a richer sequential grammar than the transition matrices reveal alone. The depth of this memory, indexed by the rate of decline in *F_N_* , tracks state-space complexity across animals: FVB16, with the smallest alphabet (*k* = 7), shows the steepest proportional drop, while FVB18, with the largest seizure count and *k* = 8, shows the highest absolute *F*_0_ values (≈ 2.3 bits), reflecting its greater marginal burst-type diversity.

**Figure 9.**
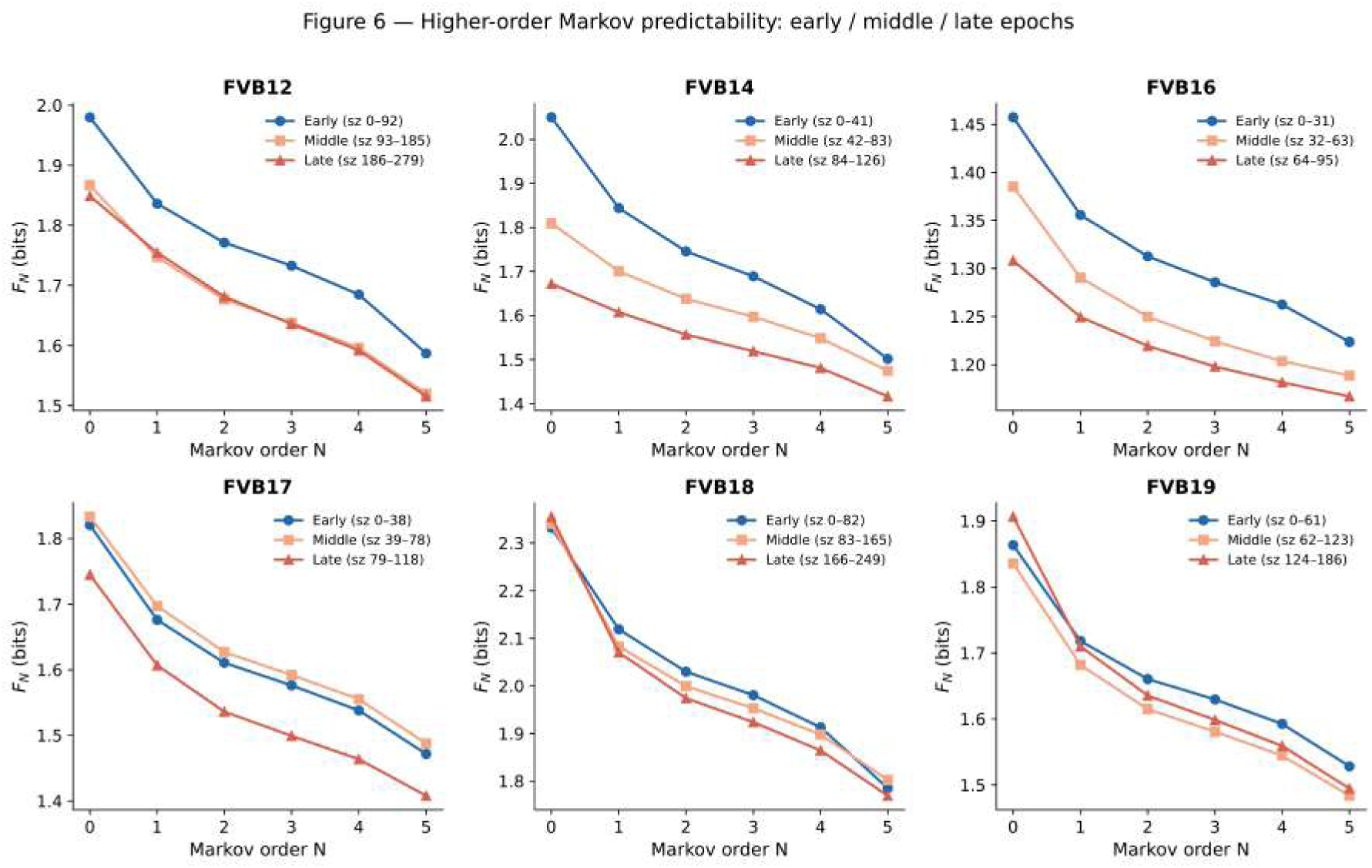
Burst sequences carry higher-order Markov memory that is non-stationary across the chronic recording. Block-entropy rate increment *F_N_* (bits; Equation 7) as a function of history length *N* (0 ≤ *N* ≤ 5) for the early (blue; first third of seizures), middle (orange; second third), and late (red; final third) epochs of each animal’s record. A flat *F_N_* for *N* ≥ 1 would indicate a first-order Markov process; a monotone decline indicates higher-order memory. In all six animals *F_N_* declines from *N* = 0 to *N* = 5: burst sequences are not first-order Markov and carry memory extending at least three to five burst states into the past. Rate of decline and absolute values track state-alphabet size: FVB16 (*k* = 7) shows the steepest proportional drop; FVB18 (*k* = 8) the highest *F*_0_ (≈ 2.3 bits). Epoch curves are non-superimposed: in FVB12, FVB14, and FVB17, the late epoch shows lower *F_N_* at all *N* than the early epoch, indicating that both burst-type diversity and higher-order memory depth decrease as the disease matures. FVB18 shows the opposite ordering for intermediate *N* ; FVB19 shows an early/late curve crossing, reflecting non-monotone reorganisation of the temporal grammar.

The temporal evolution of this higher-order structure is equally revealing. In FVB12, FVB14, and FVB17, the late-epoch *F_N_* curve lies consistently below the early-epoch curve at all values of *N* : both the marginal diversity of burst states (captured by *F*_0_) and the effective memory depth (captured by the slope for *N* ≥ 1) decrease as the disease progresses. The network, in other words, not only becomes more stereotyped in which burst states it visits, but also loses memory of its own recent trajectory. This finding mechanistically extends and deepens the stabilisation signal seen in the Frobenius and KL analyses: as the epileptic attractor consolidates, the dynamical grammar becomes simultaneously sparser, less diverse, and shorter-ranged in its temporal dependencies, as though the seizure is increasingly channeled through a narrow, well-worn path in state space.

However, not all animals follow this trajectory. FVB18 shows a non-monotone epoch ordering, with the middle epoch exhibiting the highest *F_N_* , while FVB19 shows a crossing of early and late curves, indicating a more complex, non-monotone reorganisation of the burst grammar over time. These individual differences reinforce the conclusion drawn from the Frobenius and KL analyses that the pace and form of grammatical consolidation are heterogeneous across animals, and may reflect genuine biological differences in the timing and completeness of post-SE network reorganisation.

### 20. A shared burst-state vocabulary reveals progressive individualisation of ictal dynamics across the cohort

All analyses above share a representational limitation: because *k*-means is fitted independently per animal, burst-type labels are arbitrary and cannot be directly compared across individuals. To move toward a population-level view, we pooled all *N* ≈ 1.9 × 10^6^ burst events from the six animals, standardised morphological features across the cohort, and re-clustered with a common *k*-means model (Supplementary Figure S3). The elbow criterion selected *k*^∗^ = 6 shared burst states. UMAP visualisation confirms that these six states occupy coherent, well-separated manifolds in morphological feature space (Supplementary Figure S3A), and, critically, bursts from all six animals contribute to every cluster region (Supplementary Figure S3B), demonstrating that the morphological vocabulary of burst types is broadly shared across the cohort despite differences in the per-animal *k* values (6–9).

Re-estimating per-animal Markov matrices in this common six-state space (Supplementary Figure S3C,D) confirms that transition topologies remain sparse and animal-specific even when the state vocabulary is held constant: preferred transition hubs differ across animals, and FVB14 emerges as the most idiosyncratic individual, with the largest pairwise Frobenius distances to all other animals.

The most striking result from the common-space analysis is a sign reversal in the temporal Frobenius trend. Whereas the per-animal LME yielded a significant negative slope (*β*^^^ = −0.078, *p* = 0.026; disjoint-window design, Methods), the equivalent model in the common state space yields a significant *positive* slope (*β*^^^_window_ = +0.30, SE = 0.13, *p* = 0.019), with predominantly positive Kendall *τ* values at the individual level (FVB12: +0.21; FVB14: +0.40; FVB18: +0.45; FVB19: +0.57; Supplementary Figure S3E). Both directions are therefore independently statistically significant, sharpening rather than merely suggesting the reversal.

When each animal uses its own optimised state labels, the per-animal clustering absorbs some of the epoch-to-epoch variability into the state definitions themselves, thereby partially masking how the network navigates the shared morphological space over time. When a universal vocabulary is imposed, a different dynamic becomes visible: as epilepsy matures, each animal’s pattern of burst-type usage drifts in its own particular direction through the common state space, so that transition matrices projected onto shared axes progressively diverge from each other. In other words, within each animal the grammar consolidates toward local stability, but across the cohort, animals become increasingly individualised in how they traverse the shared burst-type vocabulary. This sign reversal is not a methodological inconsistency but a substantive finding: it reveals that the temporal direction of seizure-grammar change is sensitive to the representational frame of reference, and it suggests that epileptic network maturation drives not only local stereotyping but also inter-individual divergence in ictal dynamics.

### 21. Burst-type usage profiles carry a reproducible animal-identity signature

The pooled analysis above shows that animals share a common vocabulary of six burst types but arrange them into particular transition structures (Supplementary Figure S3C,D). We next asked whether this heterogeneity is expressed strongly enough to be detected from a single seizure, i.e. whether the burst-type composition of one seizure alone is a reproducible fingerprint of the animal it came from.

Using the same *k*^∗^ = 6 pooled burst types, we represented every seizure (*N* = 1,059 across the cohort; Table I) as a burst-type usage profile – the normalised frequency of each of the six types among that seizure’s bursts – and trained a random-forest classifier (300 trees, balanced class weights) to predict the source animal from this profile alone, using stratified 5-fold cross-validation. Out-of-fold predictions reached a balanced accuracy of 0.60, more than three and a half times the chance level of 1*/*6 ≈ 0.17 (Supplementary Figure S4A), showing that burst-type composition alone identifies the animal of origin well above chance.

Classification accuracy was, however, far from uniform across animals (Supplementary Figure S4B): FVB12 (0.85) and FVB18 (0.75) were reliably identified, whereas FVB16 (0.47), FVB17 (0.38) and FVB19 (0.56) were markedly harder to distinguish, echoing the graded rather than categorical individual differences already apparent in the transition-matrix analyses.

Feature-importance ranking of the classifier singled out burst types 3, 1 and 5 as the most discriminative (importances of 0.21, 0.18 and 0.17, versus 0.16, 0.15 and 0.13 for the remaining three types), although the spread across all six types was modest, indicating that animal identity is carried by the overall usage vector rather than by one or two particular types. These three types corresponded to comparatively rare, morphologically extreme events: type 1 was the rarest (1.4% of bursts) and most extreme, combining the longest duration (291 ms), largest arc-length (13,777) and highest peak count (16 peaks/burst) of any cluster; type 3 combined the largest amplitude excursions (peak 409, trough −766) with moderate duration (84 ms); type 5 was of intermediate duration (101 ms) and the highest oscillation frequency among the top three (137 Hz).

These three types also differed systematically in their placement within the seizure (Supplementary Figure S4C): type 3 occurred early and with a tight latency distribution (median 21 s post-onset, IQR 14–30 s), type 1 slightly later with a similarly narrow spread (median 25 s, IQR 18–34 s), while type 5 occurred substantially later and with a much broader spread (median 54 s, IQR 12–80 s). Their circadian occurrence (Supplementary Figure S4D–F), in contrast, was similar across the three types, all peaking during daytime hours (06:00–16:00 UTC), matching the overall circadian seizure pattern already reported in Supplementary Figure S2 rather than showing type-specific phase preferences.

## III. DISCUSSION

Our results show that ictal LFP dynamics in the pilocarpine FVB mouse model are organised by a structured burst grammar rather than by random state sampling. Across animals, burst types formed sparse, animal-specific first-order transition matrices, with each state tending to transition to only a limited subset of successors. This supports the view that seizures unfold along constrained dynamical pathways shaped by the epileptic network architecture, rather than as undifferentiated oscillatory episodes ([6, 33]).

A key finding is that seizure sequences carry memory beyond first-order Markov statistics. The block-entropy analysis revealed that longer burst histories improve prediction of the next state in every animal, demonstrating that the symbolic seizure stream contains higher-order temporal dependencies. Consequently, the seizure grammar cannot be fully captured by pairwise transition statistics alone; the internal organisation of ictal activity depends on a broader sequence context. Such structure is precisely what symbolic dynamics and information-theoretic measures are designed to reveal ([6]).

We also observed that seizure microstructure changes over the chronic recording period. This is evident both for burst-type *occupancy* and, once analysed with a sliding window, appropriately masked for under-sampled transition-matrix rows, and tested with an autocorrelation-corrected statistic (Methods; Results), for transition *topology* : the symmetric Kullback–Leibler divergence between successive seizure occupancy distributions was consistently largest within the first ∼ 20 seizures of the chronic phase and then declined, and the Frobenius distance between successive transition matrices showed a significant negative trend at the group level (*β*^^^ = −0.018, *p* = 1.3 × 10^−4^), consistent in direction and significance with the supplementary disjoint-window analysis (*β*^^^ = −0.078, *p* = 0.026). This pattern is consistent with progressive network reorganisation following status epilepticus, in which an initially unstable ictal grammar gradually consolidates into a more stereotyped, attractor-like regime, though the effect size for transition topology is modest relative to between- and within-animal variability (Results), so our data support the consolidation account most clearly for *which* burst types are used, with the precise rules governing their succession changing in the same direction but less dramatically. The three-stage evolution of the pilocarpine model, from status epilepticus through a seizure-free latent period to chronic spontaneous seizures, is well established [33], and the temporal stabilisation of burst transition statistics we report here parallels this macroscopic disease trajectory, evident in both its occupancy- and topology-based components.

Circadian phase also appeared to modulate seizure microstructure in a subset of animals. Normalised burst-state entropy differed significantly across time-of-day bins in two of six mice, indicating that seizures occurring at different circadian phases may recruit distinct internal burst repertoires. This extends the known circadian modulation of seizure timing to the level of within-seizure organisation. Prior work has established that seizure occurrence follows circadian and multiday rhythms across species and epilepsy syndromes [1, 20, 27], and that the mechanisms underlying these rhythms likely involve an interplay between the circadian clock and sleep–wake processes [30]. Our data suggest that circadian influence can reach beyond the probability of seizure onset to modulate the dynamics of seizure unfolding itself.

The finding that a single seizure’s burst-type usage profile predicts the source animal with balanced accuracy 0.60 (chance = 0.17; Supplementary Figure S4A) provides converging, independent evidence for the individualisation already implied by the sign reversal of the pooled Frobenius/LME analysis (Section on shared burst-state vocabulary): animals do not merely share a common vocabulary of burst types that they arrange into particular transition sequences, but also differ systematically in how frequently each type is expressed within a given seizure, and this signature is stable enough to be read out from single seizures rather than only from long aggregated sequences.

Two observations temper a purely biological reading of this result. First, per-animal accuracy correlated strongly with the number of seizures available for that animal (Spearman *ρ* = 0.89, *p* = 0.019, *n* = 6; Table I), so part of the spread in Supplementary Figure S4B likely reflects the statistical power of the cross-validation rather than a graded biological distinctiveness between, say, FVB12 and FVB17. Second, because burst-type usage is derived from LFP morphology, systematic differences between animals could in principle stem from electrode placement, impedance drift, or other session-specific recording factors rather than physiologically meaningful individual variability; our single-electrode design cannot fully rule this out, and repeated recordings under matched hardware conditions would be needed to do so directly.

We note, however, that the three burst types driving classification also carry a distinctive and reproducible *within-seizure* timing signature (type 3 and type 1 cluster tightly in the first 20–35 s post-onset, whereas type 5 is both later and far more dispersed; Supplementary Figure S4C), a pattern referenced to a physiological landmark (seizure onset) that a purely instrumental artefact would not be expected to reproduce. Their circadian occurrence, by contrast, tracked the cohort-wide day/night seizure pattern rather than differing between types (Supplementary Figure S4D–F), indicating that the classifier is exploiting consistent within-seizure dynamics rather than a circadian confound. Taken together, we interpret the animal-identity signal as most consistent with genuine, if graded and unevenly powered, individual differences in burst-type deployment, adding a single-seizure line of evidence to the broader picture of progressive individualisation of ictal dynamics developed elsewhere in this study – while flagging replication in a larger, multi-electrode cohort as the appropriate next test of this interpretation.

The present study also has methodological implications. Encoding seizures as symbolic sequences makes it possible to quantify internal ictal structure in a compact and interpretable way, while preserving enough information to detect non-stationarity and higher-order memory. This approach is conceptually aligned with previous work demonstrating that epilepsy alters information-processing complexity in neural circuits [6]. The animal-specific clustering strategy used here means that absolute state labels are not directly comparable across subjects; what matters biologically is the reproducible existence of structured burst repertoires and their transition rules within each animal. However, our study presents some limitations. The cohort is small, includes only male mice, and chronic recordings do not sample all stages of epileptogenesis uniformly across animals. The circadian effects are therefore best interpreted as proof of principle rather than as a universal property of the model. Furthermore, single-channel LFP recording limits mechanistic inference about the network generators of individual burst states. In particular, seizure onset in the pilocarpine model need not originate from a single, fixed focus, and our single-electrode design cannot localise the site or extent of ictal onset for any given seizure. Consequently, part of the between-animal variability in burst-type composition and transition structure reported here – and potentially part of the within-animal evolution of that structure across the chronic recording – may reflect differences, or changes over time, in the anatomical origin or spatial extent of the epileptic network generating each seizure, rather than purely a change in the dynamical stereotypy of a fixed generator. Multi-electrode recordings spanning candidate seizure-onset regions would be needed to disambiguate these possibilities. Nonetheless, the convergent evidence from sparse transition matrices, non-stationary dynamics, higher-order memory, and circadian modulation strongly supports the conclusion that seizures contain a meaningful internal grammar.

More broadly, these findings suggest that symbolic dynamical analyses can bridge seizure phenomenology and network-level disease evolution. Aligning state spaces across animals and extending recordings to span the full latent-to-chronic transition may help determine whether the observed grammar reflects a common epileptic motif or a family of subject-specific attractor trajectories. Within the present data, however, the central message is clear: individual seizures are structured dynamical events, and that structure evolves with disease progression and is modulated by circadian phase [1, 6, 33].

## DATA AND CODE AVAILABILITY

Raw EEG data are available from the corresponding author upon reasonable request. The burst detection and symbolic analysis code is available at github.com/ViniciusLima94/epiburstdetection.

## AUTHOR CONTRIBUTIONS

AG and CB acquired the data; VL and DD conceived the research; VL performed the analysis and prepared the figures; VL and DD wrote the first version of the manuscript; VL, AG, VJ, CB, and DD edited and revised the manuscript.

## ACKNOWLEDGEMENTS

This research has received funding from the European Union’s Horizon Europe Programme under the Specific Grant Agreement No. 101147319 (EBRAINS 2.0 Project). It has also received funding from the European Union’s Horizon Europe Programme under the Specific Grant Agreement No. 101137289 (Virtual Brain Twin Project).

---

**Figure S1.**
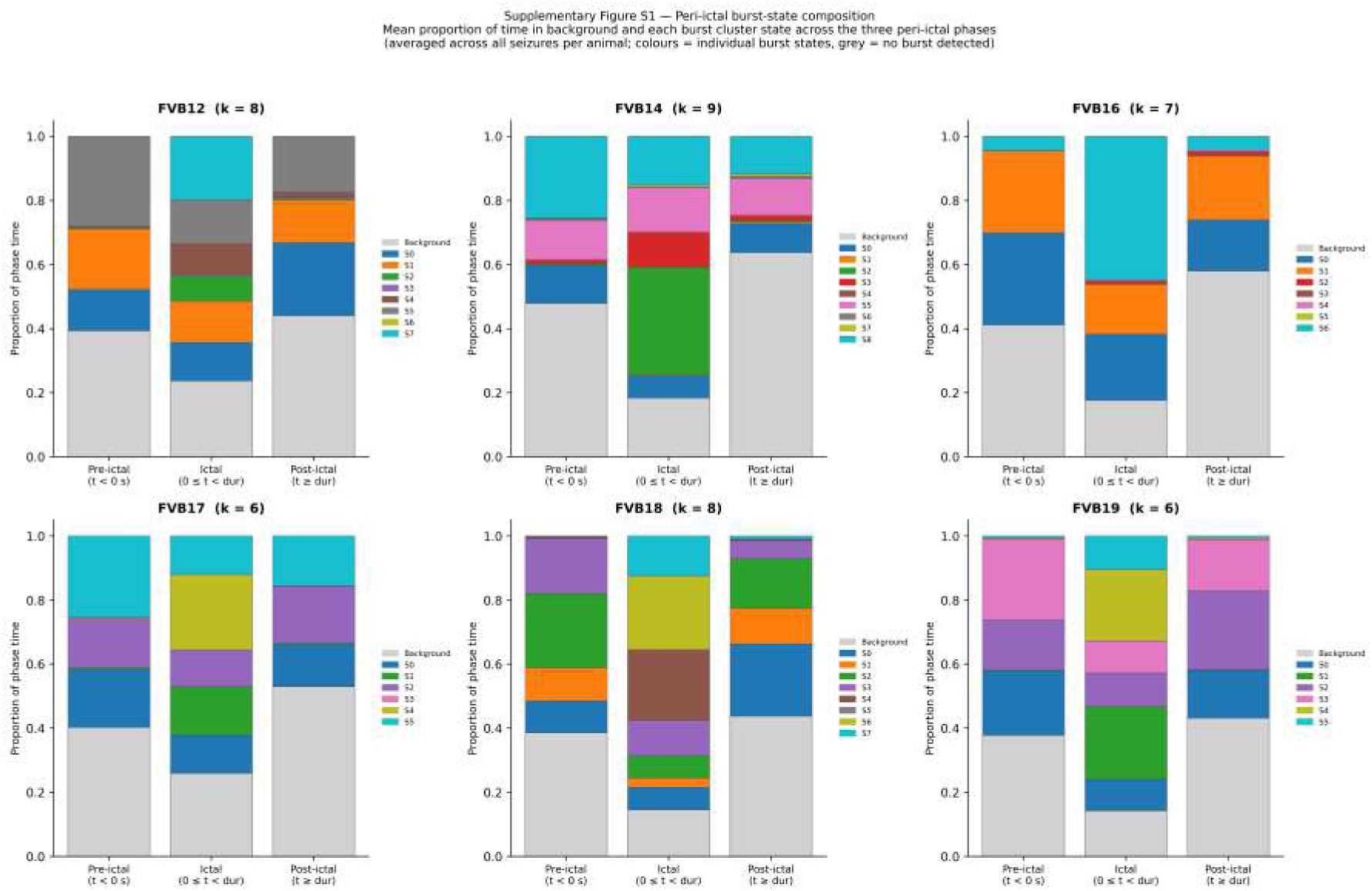
Peri-ictal burst-state composition (Supplementary Figure S1). For each animal (panels), stacked bar charts show the mean proportion of phase time occupied by background activity (grey) and each burst cluster state (colours; tab10 palette, same *k* as the per-animal analysis) across three periictal epochs: pre-ictal (*t <* 0 s, i.e. the 30 s baseline window before seizure onset), ictal (0 ≤ *t < d*_seiz_, where *d*_seiz_ is the actual seizure duration derived from EDF annotations), and post-ictal (*t* ≥ *d*_seiz_, i.e. the post-discharge window). Proportions are averaged across all seizures for each animal. In all animals, background (grey) dominates the pre-ictal phase, and the fraction of time in burst states rises sharply at seizure onset and remains elevated throughout the ictal window. The composition of burst states shifts between the ictal and post-ictal epochs in most animals: states that are rare during background activity are preferentially recruited during the sustained discharge, while the post-ictal window shows partial return toward the pre-ictal composition. The magnitude of this peri-ictal state-composition shift differs across animals, reflecting heterogeneous seizure phenotypes within the cohort.

**Figure S2.**
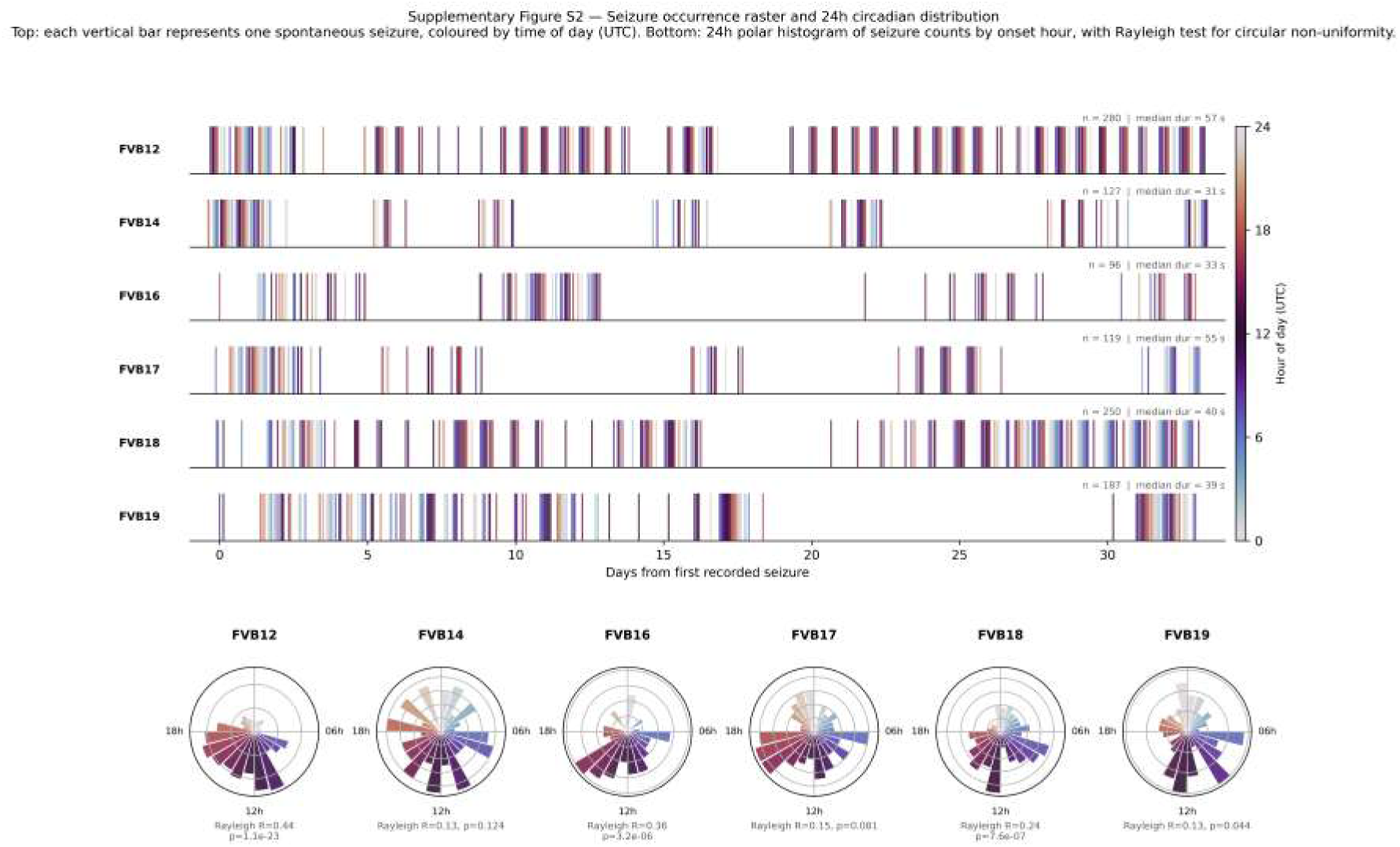
Seizure occurrence raster and 24h circadian distribution (Supplementary Figure S2). Top: Each row corresponds to one animal. Every vertical bar marks a spontaneous seizure onset; the x-axis shows days elapsed since the first recorded seizure of that animal, placing all recordings in a common relative timeline. Bar colour encodes the UTC hour-of-day of each seizure using the twilight cyclic colourmap (dark purple ≈ midnight, light yellow ≈ noon), making circadian clustering directly visible as colour banding within each row. The total seizure count and median duration are annotated per row. Several observations emerge: (i) recording durations range from ∼35 days (FVB16, *n* = 96) to ∼75 days (FVB12, *n* = 280); (ii) most animals show non-uniform temporal clustering with apparent high-density and low-density periods, suggesting multi-day cycles of seizure propensity [2]; (iii) colour banding (e.g. FVB18, FVB19) reveals circadian preferential timing, with seizures clustering in certain time-of-day windows, consistent with the entropy analysis in Figure 7. **Bottom:** 24-hour polar (rose) histogram of seizure onset hour per animal (24 one-hour bins; 0 h at top, clockwise), with the Rayleigh test for circular non-uniformity (Methods) reported below each panel. Seizure timing is significantly non-uniform in four of six animals (FVB12, *p* = 1.1×10^−23^; FVB16, *p* = 3.2 × 10^−6^; FVB18, *p* = 7.6 × 10^−7^; FVB19, *p* = 0.044) but not in FVB14 (*p* = 0.124) or FVB17 (*p* = 0.081), giving a formal statistical counterpart to the colour banding visible above.

**Figure S3.**
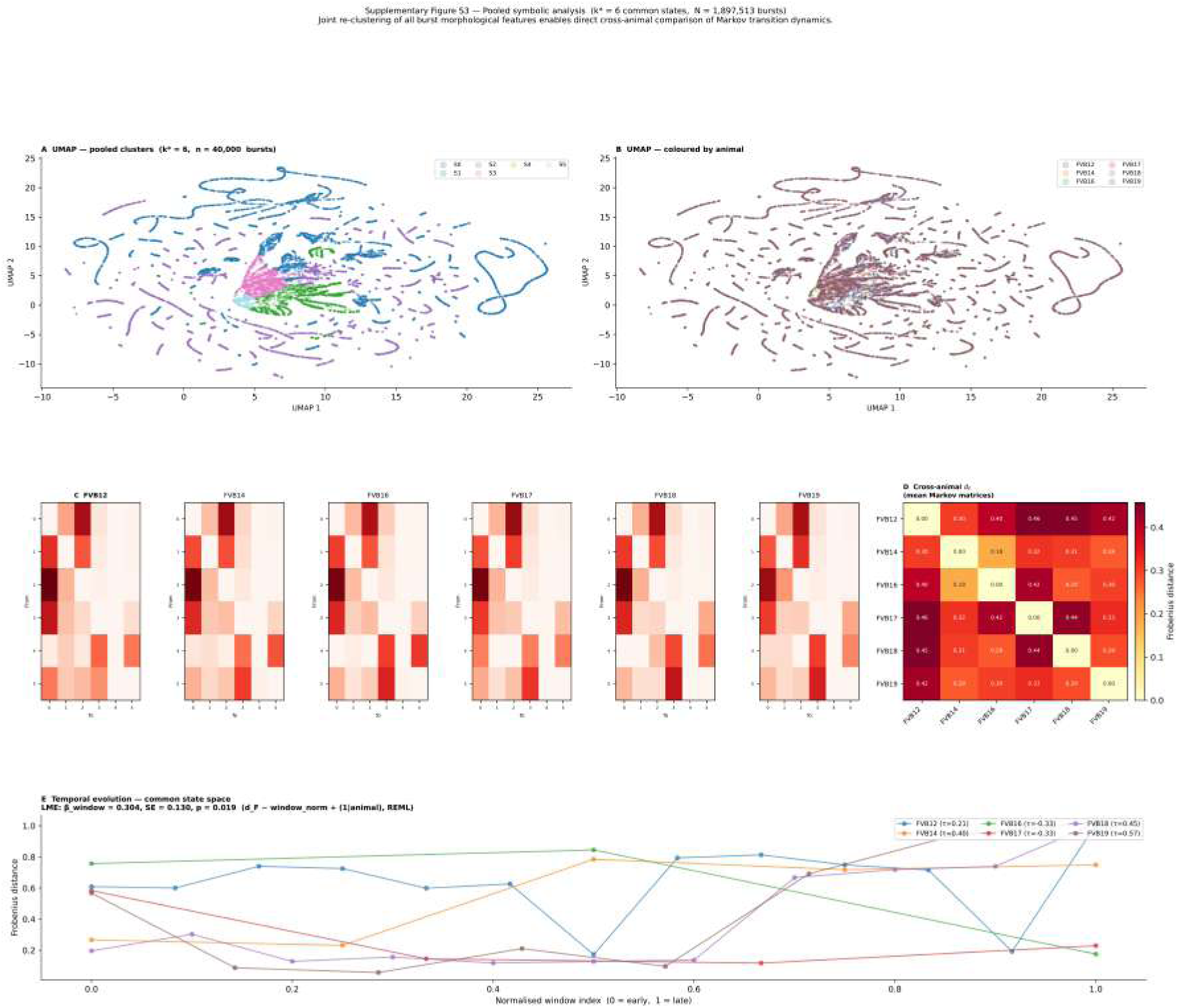
Pooled symbolic analysis in a common burst-state space (Supplementary Figure S3). All bursts from the six animals (*N* = 1,897,513 events) were jointly standardised and re-clustered with KMeans; the optimal common state count was *k*^∗^ = 6 (determined by the maximum-curvature inertia elbow on a 60,000-burst subsample). **(A)** UMAP embedding of 40,000 pooled burst feature vectors (subsampled for speed), coloured by pooled cluster assignment. Clusters form coherent, well-separated manifolds in the reduced space. **(B)** Same UMAP embedding coloured by animal of origin. Bursts from all six animals contribute to each cluster region, demonstrating that the morphological feature space is broadly shared across the cohort despite differences in per-animal *k* (6–9). **(C)** Mean Markov transition matrices per animal estimated in the common 6 × 6 state space, enabling direct cross-animal comparison of transition topology without dimensionality mismatch. Matrices remain sparse and animal-specific. **(D)** Pairwise cross-animal Frobenius distance matrix (6 × 6) computed between the mean Markov matrices in the common state space. Off-diagonal entries quantify how different each pair of animals is in their average burst-type transition topology; the diagonal is zero by construction. FVB14 shows the largest pairwise distances to most other animals, consistent with its distinctive transition hub structure (Figure 3). **(E)** Temporal evolution of the Frobenius distance between consecutive 20-seizure transition matrices in the common state space (one line per animal). The pooled LME (random intercept per animal, REML) yields *β*_window_ = +0.30 (SE = 0.13, *p* = 0.019), a *significant increase* that is opposite in sign to the per-animal result (*β* = −0.078, *p* = 0.026; disjoint-window design, Methods), revealing that the temporal direction of transition-structure change is sensitive to the choice of state-space representation (see Results for interpretation).

**Figure S4.**
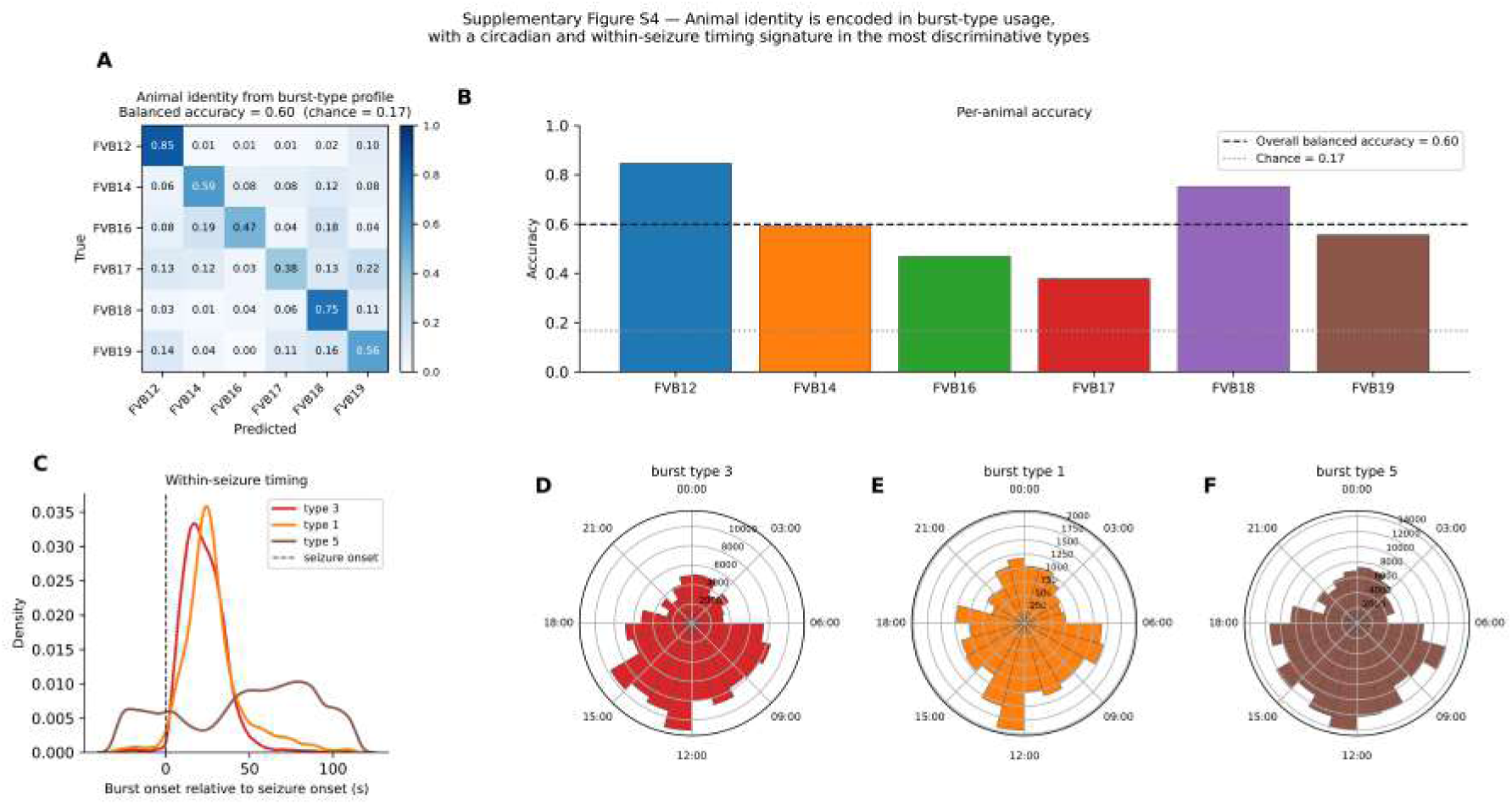
Animal classification from burst-type usage (Supplementary Figure S4). The pooled six-burst-type vocabulary (Supplementary Figure S3) was used to summarise every seizure (*N* = 1,059) as a burst-type usage profile, from which a random-forest classifier predicted the source animal (Methods). **(A)** Confusion matrix of out-of-fold predictions (5-fold stratified cross-validation), normalised by true animal. Balanced accuracy is 0.60 against a chance level of 1*/*6 ≈ 0.17. **(B)** Per-animal accuracy (confusion-matrix diagonal) as a bar chart, with the overall balanced accuracy and chance level shown as reference lines. FVB12 and FVB18 are the most reliably identified animals; FVB16, FVB17 and FVB19 are harder to distinguish. **(C)** Timing of the three most important burst types (identified by random-forest feature importance) relative to seizure onset. Type 3 and type 1 cluster tightly soon after onset, whereas type 5 occurs later and with a broader spread. **(D–F)** Time-of-day distribution (polar, clock-labelled) of the same three burst types. All three show a similar daytime-weighted circadian profile, consistent with the cohort-wide seizure timing pattern (Supplementary Figure S2) rather than a type-specific circadian signature.

